# Long-term Culturing of *Pseudomonas aeruginosa* in Static, Minimal Nutrient Medium Results in Increased Pyocyanin Production, Reduced Biofilm Production, and Loss of Motility

**DOI:** 10.1101/2024.02.26.582132

**Authors:** Rhiannon E. Cecil, Elana Ornelas, Anh Phan, Nahui Olin Medina-Chavez, Michael Travisano, Deborah R. Yoder-Himes

## Abstract

*Pseudomonas aeruginosa* is a multidrug-resistant opportunistic human pathogen that can survive in many natural and anthropogenic environments. It is a leading cause of morbidity in individuals with cystic fibrosis and is one of the most prevalent pathogens associated with nosocomial infections in the United States. It has been shown that this organism can survive and persist in low nutrient environments, such as sink drains. How adaptation to these types of environments influences the phenotypic traits of this organism has not been well studied. Here we implemented an experimental evolution system in which six strains of *P. aeruginosa* were subjected to low nutrient conditions over the course of 12-weeks and assessed phenotypic and genotypic changes that occurred as a result of adaptation to such environments. We observed that adaptation to low nutrient environments resulted in decreased generation time, reduced cell size, reduced biofilm formation, increased pyocyanin production, and decreased motility for some of the strains. Further, some of the evolved isolates were significantly more virulent/competitive against a phagocytic predator. This study is significant as it allows us to predict how this organism will evolve in hospital and domestic environments and can help us improve treatment options for patients.

**Importance:** Human commensal and pathogenic organisms undergo dynamic cycles across human and non-human environments. Despite the crucial implications for human health, the understanding of bacterial adaptations to these diverse environments and their subsequent impact on human-bacterial interactions remains underexplored. This study shows how *P. aeruginosa*, an opportunistic human pathogen, adapts phenotypically in response to a shift from high nutrients (like those found in the human body) to low nutrients (like those found in many other environments, like sink drains). This work also shows that, in some cases, resistance to predatory forces can evolve in the absence of a predator. This work is important as it contributes to the growing body of knowledge concerning how external, non-host-related abiotic conditions influence host-pathogen interactions.

## Introduction

Adaptations to environments with fluctuating nutrients, otherwise known as feast and famine cycles, are vitally important for microbial survival in various environmental niches including natural environments such as soil, aquatic, and the intestinal and respiratory tract of host organisms; as well as anthropogenic environments such as drains and hospital equipment (1). Adaptations employed by some microorganisms in response to such environments, such as the stringent response (1–3), cellular dormancy via reduced cell growth and the formation of spores (4, 5), and metabolic shifting (fermentation vs respiration) (6, 7), are already well-studied. Our understanding of how these adaptations evolve in opportunistic human pathogens and the implications such adaptations have on virulence potential is relatively understudied. Uncovering the mechanisms behind these adaptations holds significant potential for ecological, evolutionary, environmental, industrial, and pharmaceutical applications. For example, these insights can be leveraged to better understand microbial diversity, microbial food webs, speciation, the evolution of bacterial communities, improve bioremediation techniques for cleaning polluted environments, develop more efficient industrial processes for biofuel production, and even design new antibiotics by targeting the pathways crucial for bacterial survival under nutrient limitations.

*P. aeruginosa* is a member of the ESKAPE group of bacterial pathogens (8). It is an opportunistic pathogen and is known for its intrinsic multidrug resistance, genomic plasticity, ability to survive and form biofilms on abiotic surfaces such as hospital equipment, and ability to evade the immune systems of humans, plants, and non-human animals (9–13). *P. aeruginosa* expresses with a wide arsenal of phenotypes that serve as a means of offensive and defensive protection against competitors and predators in the environment (e.g., protozoa, nematodes) including pyocyanin production, biofilm formation, and T3SS expression (14–16). Many of these phenotypes coincidentally act as virulence factors should the bacterium encounter a human host. *P. aeruginosa* is a well-studied organism and how it adapts to host-associated niches, such as the nutrient-rich cystic fibrosis lung, has been researched in detail (17); however, how this organism evolves when forced to adapt to a low-nutrient, non-host associated environment has not been thoroughly investigated. Furthermore, comparisons between adaptations to such an environment between *P. aeruginosa* isolates from clinical origin and those from an environmental origin remains under-explored with many studies utilizing clinical isolates and excluding environmental isolates (18). Understanding how *P. aeruginosa* isolates, from clinical and environmental origin, adapt to survive in a low-nutrient medium could increase our understanding of how *P. aeruginosa* adapts in other low nutrient environments including sink drains, intubation tubes and other types of hospital equipment, human tissues, and other such reservoirs where this organism is able to thrive.

Experimental evolution provides a powerful means of understanding fundamental evolutionary processes and can also be used to increase our understanding of how virulence factors in opportunistic pathogens arise or change over time (19, 20). In this study, we used an experimental evolution system to identify how six *P. aeruginosa* strains (four clinical strains and two environmental strains) adapted to a low-nutrient, static environment over the course of multiple generations. Whether adaptations to this type of environment would result in changes to phenotypes commonly associated with growth and/or pathogenicity was then explored. We also sought to determine if long-term culture in a low nutrient environment under static conditions would result in increased or decreased fitness against a phagocytic predator, *Acanthamoeba castellanii*, which would suggest that adaptation to such environments could result in increased virulence in other organisms.

## Materials and Methods

### Bacterial strains and culture maintenance

*P. aeruginosa* strains used in this study were chosen based on their colony phenotype, pyocin profile, and source of isolation. Strain details are listed in Table 1. Cells were routinely maintained on LB (Lennox) agar plates at room temperature (22°C) or grown in LB (Lennox) broth. Prior to initiation of an experiment, a single colony would be transferred from a LB agar plate to either glass test tubes containing 5 mL of LB broth, 96-well deep plates containing 1% HL5 medium (per L: 0.5 g KH_4_PO_4_; 0.5 g Na_4_HPO_4_; 7 g yeast extract; 14 g proteose peptone, 13.5 g glucose; purchased premixed from Formedium, Catalog #HLG0102), or 96-well polystyrene plates containing 180 µL of LB broth and incubated overnight at 37°C with or without aeration as indicated. When indicated, mid-log cultures were generated from overnight liquid culture by transferring 100 µL of overnight culture to 5 mL of sterile LB broth and incubated at 37°C with aeration for 90 minutes.

**Table 1.** Strains used in study and their pyocin profile.

| Strain | Origin | Colony Phenotype | S1-type | S2-type | S1S2 immunity | R-type | F-type | Reference |
| --- | --- | --- | --- | --- | --- | --- | --- | --- |
| PA B80398 | *CF sputum | Normal Colony | 1<sup>a</sup> | 0 | 0 | 1 | 0 | (33) |
| PA B80427 | CF sputum | Small Colony | 0 | 1 | 1 | 1 | 1 | (33) |
| PA B84725 | CF sputum | Mucoid | 0 | 0 | 0 | 1 | 1 | (33) |
| PA3 | CF sputum | Normal Colony | 0 | 1 | 0 | 1 | 0 | (33) |
| SRP 17-047 | Bathroom sink drain | Normal Colony | 1 | 0 | 1 | 1 | 0 | (34-36) |
| SRP 17-055 | Kitchen sink drain | Normal Colony | 1 | 0 | 1 | 1 | 1 | (34-36) |

<sup>a</sup>1 indicates presence of the pyocin gene in the genome, 0 indicates absence from the genome.
\*CF indicates isolates taken from sputum samples from individuals with cystic fibrosis

### Experimental Evolution Methods

Each strain was cultured in 8 mL of 1% HL5 in 6-well tissue culture plates. The plates were incubated at room temperature under static conditions. Every 2 to 3 days the medium was removed with vacuum suction and replaced with 8 mL of fresh sterile 1% HL5 medium. The evolving populations were also passaged every 7 days by removing the supernatant, adding 4 mL of sterile 1% HL5, scraped with a cell scraper to dislodge the adherent cells. The suspended cells were transferred to a sterile 15 mL conical tube and centrifuged at 6,976 x *g* for 15 minutes at room temperature. The pellets were resuspended in 1 mL of 1% HL5 and 500 µl was added to 8 mL of sterile 1% HL5 in fresh 6-well tissue culture plates. A strain collection was generated at bi-weekly time points by transferring 500 µL of culture to 500 µL of sterile LB+40% glycerol (glycerol 20% final concentration) with subsequent storage at −80°C. At each dilution step, lineages were tested for contamination via colony PCR for pyocin biosynthesis genes. Pyocins are proteins exclusively produced by *P. aeruginosa.* The pyocin gene profiles were used to ensure that no cross contamination occurred between the strains throughout the experimental evolution via colony PCR using published primer sequences for the specific pyocin genes (21). Three biological replicate lineages for each strain were evolved for 12 weeks total time.

On the rare occasion that a deviation of expected pyocin profiles was observed for any given gene, the pyocin profiles were examined in the following week determine if the trend was conserved. In all cases, the pyocin profiles returned to the expected pattern. Two strains that were originally included in the dataset (not shown) were removed due to repeated loss of pyocin gene positive results, which could result from contamination or simply selection against these genes. However, we note that the final, analyzed isolates used in this study (the 12-week isolates) all matched the respective ancestral pyocin profile.

### Generation time

Overnight cultures of ancestral and evolved isolates were diluted (1:100 B83098, 17-047, 17-055 or 1:200 for B84725, PA3, and B80427) into 15 mL in 125-ml baffled flasks. Flasks were incubated with shaking at 37°C. Every hour, the O.D._600_ was measured from 1 mL of culture and plotted in GraphPad Prism v 5.04. At time points within exponential growth phase, samples were taken for survival analysis via serial dilution and drip plating on LB square petri dishes. Average bacterial concentrations were calculated for each of three biological replicates for each ancestral and evolved isolate.

### Cell Size

Overnight cultures of ancestral and evolved isolates were generated in 5 mL of LB broth in glass test tubes with aeration. 20 µL of overnight culture was transferred onto a glass microscope slide along with a drop (∼20 µL) of 1% congo red staining solution. The mixture was smeared across the slide using a coverslip then allowed to dry for 5 minutes in a biosafety cabinet. The slides were imaged under a 1000X magnification light microscope with an ocular micrometer for scale. ImageJ was used to quantify the length of cells. At least 100 cells from at least two independent experiments (n > 200 for each isolate/strain) each were measured.

### Biofilm biomass quantification

Biofilms were quantitated via a slightly modified version of a previously published protocol (22). Briefly, overnight cultures of ancestral and evolved lineages in replicates of four of each isolate were generated in 180 µL of LB in 96-well dilution plates without aeration. Biofilms were set-up by first using a 96-well transfer apparatus to transfer inocula of overnight cultures to fresh 96-well PVC plates containing 100 µL of 1% HL5. The inoculated plates were sealed with porous adhesive film (VWR, 60941-086), loosely wrapped with aluminum foil, and then placed in a humidity chamber with sterile water at the bottom. The lid of the humidity chamber was loosely sealed to allow for air flow while also maintaining a moist environment. The plates were incubated at 37°C for 48 hours or 7 days to allow for biofilm formation. Biofilms were washed two times with water then 125 µL of 0.1% crystal violet was added to each well and the plates were allowed to incubate at room temperature for 10 minutes. Afterwards, the biofilms were washed four more times with water. Then 200 µL of 30% acetic acid solution was added to each well and allowed to incubate at room temperature for 15 minutes. A 96-well plate reader was used to quantify crystal violet in solution at an optical density of 500 nm.

### Pyocyanin and pyoverdine production

Pigment levels from *P. aeruginosa* strains were determined using protocols modified from a previously published study (23). Overnight cultures were generated in 5 mL of LB broth with aeration in glass test tubes. 100 µL of overnight culture was transferred to 1 mL of LB broth in 24-well plates. The plates were sealed with porous adhesive film (VWR, 60941-086) and incubated at 37°C for 48 hours under static conditions. Post-incubation, 1 mL of each sample was transferred from the 24-well plates to 1.5 mL microfuge tubes. The tubes were then centrifuged at 21,130 x *g* for 30 minutes to pellet the bacteria, and 100 µL of supernatant from each culture was transferred to either 96-well clear plates (VWR, #10861-562) for pyocyanin quantitation or 96-well black plates (Greiner, #655077) for pyoverdine. A 96-well plate reader was used to quantify the amount of pyocyanin (absorbance at 686 nm) or pyoverdine (fluorescence 395ex/460em).

### Motility assays

The motility protocol used in this study was modified from the motility assay described in (23). Overnight cultures of ancestral and evolved isolates (each n = 3) were grown in 5 mL of LB broth in glass test tubes with aeration. Briefly, a 2 – 10 µL pipet tip was dipped in overnight culture then was used to stab approximately half-way deep into the motility agar (10 mL of 0.3% LB agar in 6-well plates). The inoculated plates were incubated at room temperature for 24 hours. The diameter of spread was measured using a metric ruler to quantify swimming distance. Isolates from each replicate lineage were tested at 12 weeks. For those that decreased, isolates were then tested at 2, 3, 5, and 7 weeks but only until the time when motility appears to be significantly reduced.

### *Acanthamoeba castellanii* culture and maintenance

*A. castellanii* strain 30010 was obtained from the ATCC and cells were routinely maintained at room temperature in T-75 tissue culture flasks containing 15 mL of 100% HL5 growth medium. The cells were passaged routinely when they reached confluence, approximately every 3 days, using a cell scraper to dislodge the cells from the bottom of the flask, then removing all but 1 mL of culture from the flask and replacing with 14 mL of sterile 100% HL5. Cell stock cultures were maintained up to 4 passages.

### Co-culture experiment with *Acanthamoeba castellanii*

After reaching confluence, approximately 6 x 10^3^ *A. castellanii* cells, based on hemocytometer counts, were transferred to wells of 6-well tissue culture plates along with 4 mL of 100% HL5 medium. The cells were incubated at room temperature for 1 hour to allow the cells time to anneal to the surface of the wells in the 6-well tissue culture plates. The 4 mL of 100% HL5 was then removed, and the wells were washed 2 times with sterile 1X PBS then 8 mL of 1% HL5 was added to the wells. Overnight cultures of ancestral *P. aeruginosa* and one of the evolved lineages of each *P. aeruginosa* strain were generated in triplicate in 5 mL of LB broth. Mid-log cultures were generated as previously described above until an O.D._600_ between 0.8 and 1.0 was reached for all strains. *P. aeruginosa* cells were added to the 6-well tissue culture plates in a ratio of 1 *P. aeruginosa* cell to 10 *A. castellanii* cells in 3 biological replicates per *P. aeruginosa* isolate.

The plates were incubated at room temperature for 16 days. Every three days the medium was replaced by using vacuum suction and replaced with 8 mL of sterile 1% HL5. *A. castellanii* cells were enumerated on days 0, 3, 6, 8, 11, 13, and 16 via direct cell count at 400X magnification. *P. aeruginosa* cells were enumerated on day 16 via serial dilution and plating on LB agar plates.

### Genome sequencing

One mL of each ancestral and one of the evolved lineages for B80398, B80427, 17-047, and 17-055 was grown in LB medium and genomic DNA was isolated using a commercial kit (Promega Wizard Genomic DNA Purification Kit) according to the manufacturer’s instructions. DNA was shipped on dry ice to Novogene for genome sequencing. A total amount of 0.2 μg DNA per sample was used as input material for the DNA library preparations. Briefly, genomic DNA sample was fragmented by sonication to a size of 350 bp. Then DNA fragments were endpolished, A-tailed, and ligated with the full-length adapter for Illumina sequencing, followed by further size selection and PCR amplification. After PCR products were purified by AMPure XP system (Beverly, USA). Subsequently, library quality was assessed on the Agilent 5400 system (Agilent, USA) and quantified by QPCR (1.5 nM). The qualified libraries were pooled and sequenced on Illumina NovaSeq platforms with PE150 strategy at ∼100X coverage, according to effective library concentration and data amount required.

### Variant Calling Analysis

To compare ancestral and evolved lineages, and measure punctual changes related to adaptation, we performed variant calling analysis. After sequencing, raw read quality control was performed using FastQC v.0.20.0 (24) to identify adapters and then remove them using Cutadapt v4.2 (25). We then proceed to trim sequences with low quality bases using Trimmomatic (26) with the following parameters LEADING:10 TRAILING:3 SLIDINGWINDOW:4:30 MINLEN:35. Paired-end high quality reads were mapped to the reference genome, *P. aeruginosa* UCBPP-PA14, using Burrows-Wheeler Aligner (BWA) v.0.7.17 (27). Post-alignment processing, including sorting, indexing, and duplicate removal, was conducted with Samtools (28) to ensure clean and accurate input data for variant analysis. Base recalibration and indel realignment were performed using GATK’s Best Practices workflow to correct systematic errors introduced during sequencing (29–31). Variants were called using GATK’s HaplotypeCaller, configured to detect both SNPs and indels across the entire genome. Stringent filtering criteria were applied to identify high-confidence variants, including minimum mapping quality scores, read depth thresholds, and variant quality scores. SnpEff (32) was used to identify the genes where the variants were located or the closest genes to a variant and to predict the effects and impacts of the variants on the genes and protein products.

### Data Availability

Raw sequencing data can be found in NCBI at the following BioProject ID: PRJNA1194290

### Statistics

Data was analyzed with one-way ANOVA with Tukey’s or Dunnett’s post-test or with t-tests as described below in GraphPad Prism 5.0.4 as indicated. Spearman correlations were calculated in GraphPad Prism 9.0.

## Results

To test how a variety of *P. aeruginosa* phenotypes change in response to forced growth in a low-nutrient environment, we conducted a 12-week-long experimental evolution on six *P. aeruginosa* strains. Four of these strains were of clinical origin (PA B80398, PA B80427, PA B84725, and PA3) and two strains were of environmental origin (SRP 17-047 and SRP 17-055) as they were isolated from household sink drains. These strains were chosen based on their colony phenotypes, pigment production, and pyocin profiles and these strains represent a wide range of *P. aeruginosa* morphotypes. Four of the strains (PA B80398, PA3, SRP 17-047, and SRP 17-055) present with “normal” colony morphology. The normal colony phenotype of *P. aeruginosa* is frequently isolated from acute infections or environmental samples and is associated with more aggressive virulence factors such as pyocyin production and T3SS as well as increased motility and a predisposition to planktonic lifestyle (37–39). The other two strains, PA B80427 and PA B84725, are small colony and mucoid respectively, both of which are often isolated from chronic infections and are not known to be isolated from environmental samples. The small colony and mucoid phenotypes commonly present with defensive virulence factors to avoid detection by the host’s immune system such as reduction in motility and increased biofilm formation (38, 40). The mucoid phenotype also overproduces the carbohydrate polymer alginate making them difficult to phagocytose (41).

An experimental evolution was designed to mimic the conditions that *P. aeruginosa* would encounter in a non-host environment, such as the soil or an abiotic surface such as a sink drain. To mimic these environments, *P. aeruginosa* was cultured over time under static conditions (to encourage niche specialization) in a low-nutrient medium (1% HL5 with glucose), which contains dilute yeast extract and peptone along with sodium and potassium buffers. This medium was chosen as *P. aeruginosa* readily utilizes glucose and amino acids as carbon sources, amino acids stimulate biofilm formation and swarming activities, and for subsequent amoebae studies (33, 42–44). Every two days, the medium was removed via vacuum suction [simulating the flow of liquid down a sink drain (i.e. frequent mechanical removal)]. This removes most of the planktonic cells, thus selecting for cells that can attach to the surface of the wells. Every 7 days the medium was removed, and the adherent cells were scraped from the wells and diluted back into sterile 6-well tissue culture plates containing sterile 1% HL5. This process was continued for 12 weeks generating three biological replicate lineages of the six *P. aeruginosa* strains. Interestingly, the normal and mucoid colony strains remained the same in terms of colony morphology over the course of the experiment. However, all three replicate lineages of the small colony variant, B80427, underwent a morphological shift to a normal colony phenotype during the evolution experiment. All the starting ancestral isolates and isolates from the end of the evolution experiment (i.e. 12-week evolved isolates) were tested for a variety of phenotypes associated with growth, long-term survival, or virulence.

### Overall cellular changes during evolution in a low nutrient medium

During evolution experiments, an increase in fitness of the evolved lineage compared to its ancestor usually occurs as the organisms adapt to their conditions. One global indicator of fitness in bacteria is generation time. A decreased generation time in 1% HL5 at room temperature would indicate that the evolved isolates have an increased fitness in this environment compared to their ancestors and thus this measurement serves as a control for the experimental evolution. We assessed growth of all ancestral and representative isolates from evolved lines in 1% HL5 over time and calculated the generation times for each isolate during exponential phase growth. Both generation times and growth curves in 1% HL5 were used to assess overall growth patterns at two temperatures, 22°C which is the temperature used for the experimental evolution and at 37°C, which is the optimal growth temperature for most *P. aeruginosa* strains. At 22°C, generation times were on the orders of hours, not minutes, due to the low nutrient conditions as well as the temperature. Most evolved isolates from three strains, B80427, PA3, and 17-047, showed a decreased generation time compared to their associated ancestral strain. This suggests that they adapted to the conditions of the experimental evolution. Two individual isolates from the other strains, in B80398 and B84725, showed an increase in generation times while the other two evolved isolates did not significantly change compared to their ancestral parent. One strain, 17-055 did not show a change in generation times for any of the evolved isolates.

We then assessed whether evolving in 1% HL5 for 12 weeks at room temperature also conveyed an advantage at a higher temperature due to simply the medium alone. We calculated the generation time for all strains grown in 1% HL5 at 37°C, generally considered the optimal temperature for *P. aeruginosa* (though we note that we did not identify the true optimal temperatures for all strains in this study). Three strains, B84725, PA3, and 17-055, have the majority of evolved isolates showing a reduced generation time while almost all of the others are not statistically different than their respective ancestors. Comparing the two temperature studies, we also observed that *P. aeruginosa* B3098 evolved line 2 showed an increase in generation time (i.e. slower growth) at both 22° and 37°, suggesting that this isolate did not adapt to the low nutrient conditions of the experimental evolution for some reason. Interestingly, the B84725 showed a different pattern between the two temperatures with an increased generation time at 22°C and a slightly decreased generation time at 37°C. This particular strain is the only mucoid isolate in our strain panel and it has previously been shown that the expression of alginate biosynthesis genes is higher at 20°C than 37°C (45).

By examining the growth curves of each lineage, we observed that there were many significant differences between evolved and ancestral lineages at 22°C but not at 37°C based on repeated measures one-way ANOVAs (Fig. S1). Taken all the growth metrics together, this suggests that many of the genetic changes that occurred during the evolution served to optimize growth at room temperature and some of these changes may have additional benefits, even at higher temperatures.

Cell size is also linked to metabolic fitness as an increased surface volume ratio results in amplified nutrient diffusion into the cell resulting in increased metabolic efficiency, especially in low nutrient conditions (46, 47). We hypothesized that an overall decrease in cell size would be observed in the evolved isolates as the low-nutrient concentration of the growth medium would encourage a smaller surface area to volume ratio. Many of the evolved isolates appeared to have reduced cell length compared to the ancestor at 22°C (Fig. 2). A few evolved isolates that increased in cell length at this temperature, including B80427 evolved lineage 1 (E1) and B84725 evolved lineage 3 (E3). Curiously, the evolved lineages of PA3 all increased in size compared to the ancestor; however, this ancestral strain was relatively small to begin with, which might imply that PA3 was already adapted to lower nutrient conditions. At 37°C, most of the differences in cell length were not observed. However, all evolved lineages of B84725 showed a decrease in cell length while a few evolved isolates, PA3 E3, 17-047 E2 and E3, 17-055 E2, had a modest increase in cell length. Our cell size data overall suggests that a small cell size was not strongly selected for but that individual lineages had slight increases or decreases in cell length. This data is confirmed when aggregating the data from the evolved lineages as well (Fig. S2).

**Figure 1.**
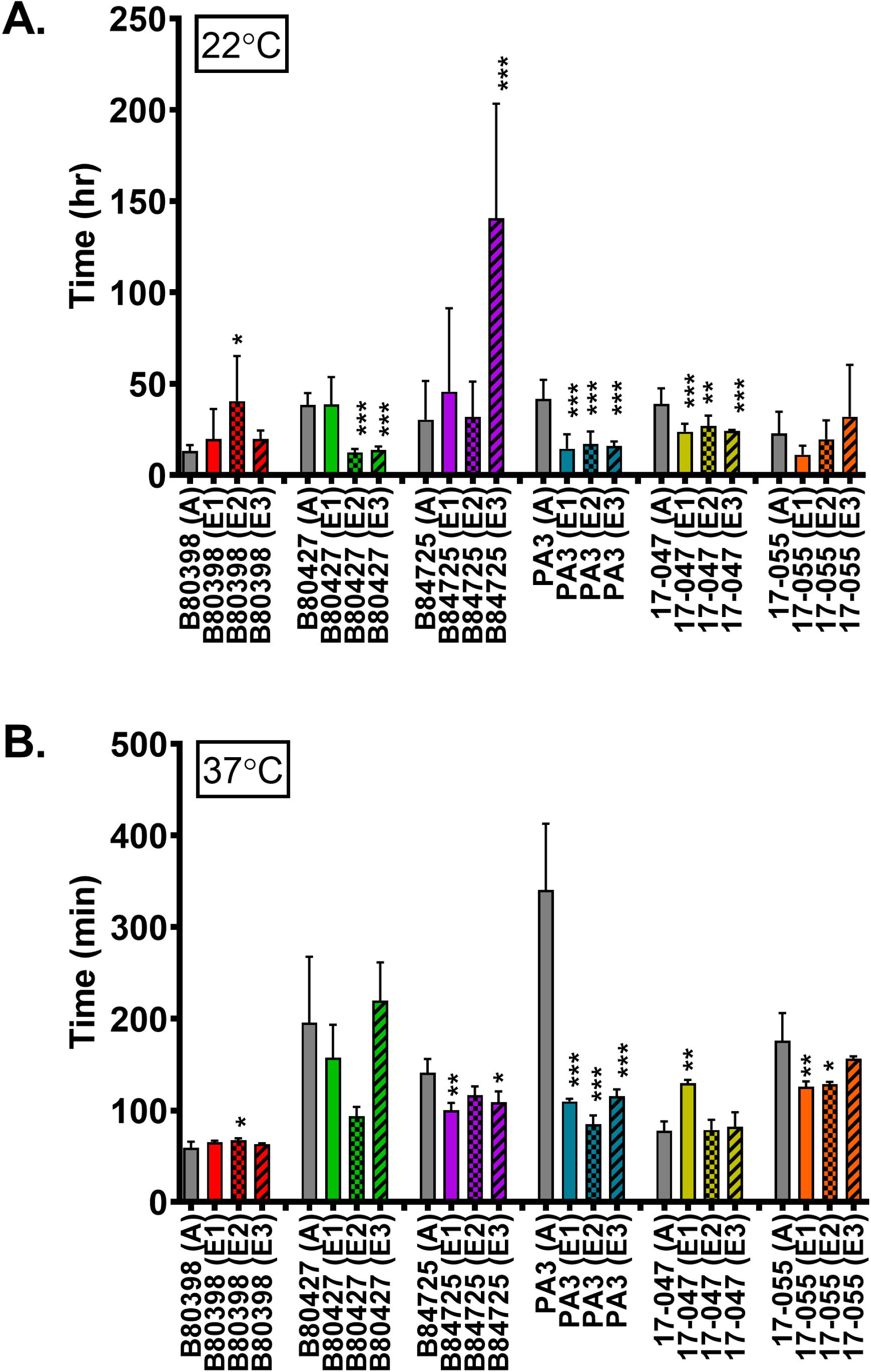
Relative generation time of ancestral and evolved *P. aeruginosa* isolates in 1% HL5. Generation times were calculated for each strain growing at 22°C (panel A) or 37°C (panel B). Grey bars represent the ancestral strains and colored bars represent the average of three replicates of each evolved lineage. Data analyzed with one-way ANOVA with Dunnett’s post-test comparing all evolved lineages to the ancestor. Error bars represent standard deviation. * p<0.05, ** p<0.01, *** p<0.001.

**Figure 2.**
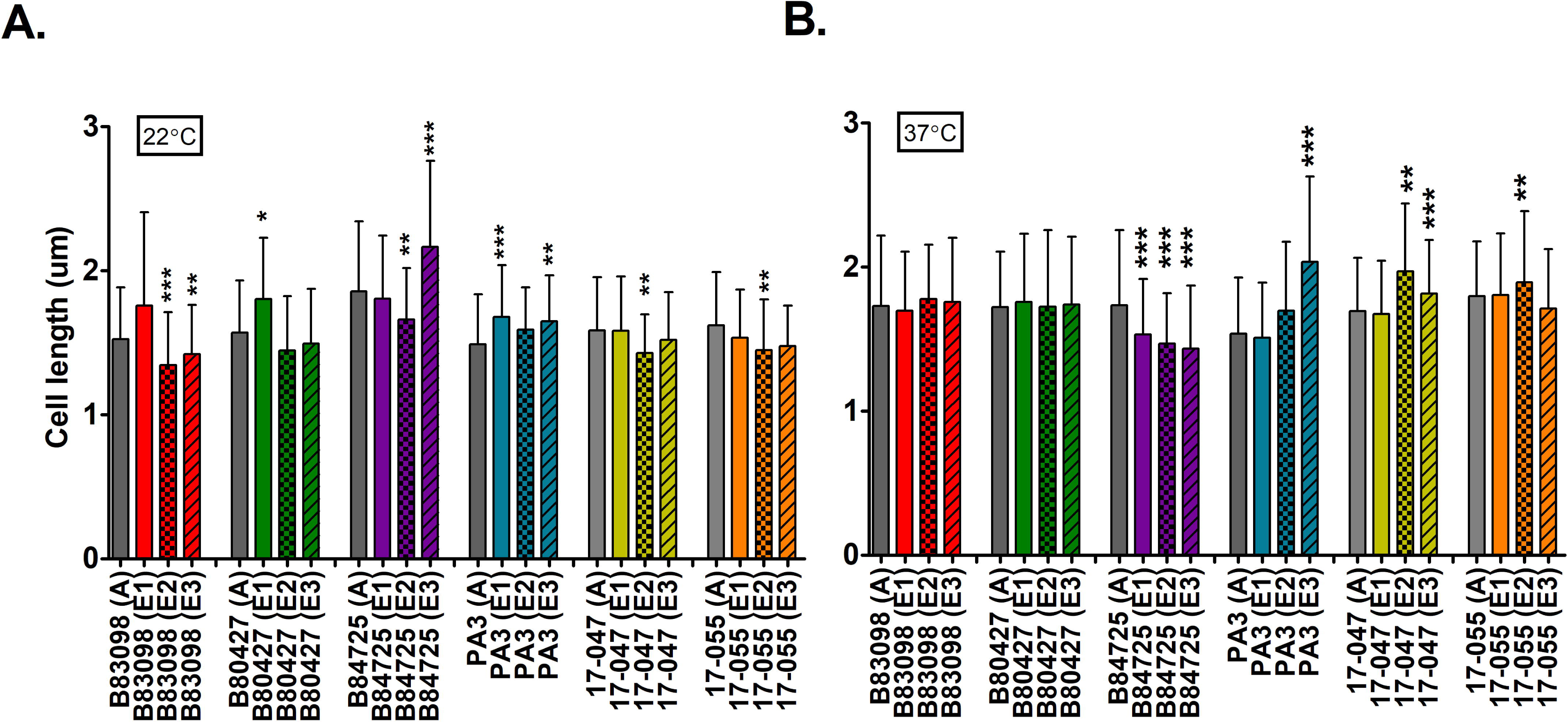
Cell size of ancestral and evolved lineages grown in 1%HL5 medium. Panels show cell sizes for strains growing at (A) 22°C or (B) 37°C. Grey bars represent average values for the ancestor strain while colored bars represent the average of three evolved isolates. An (A) in the x-axis indicates the ancestral isolate, and (E) represents evolved isolates. Data analyzed with one-way ANOVA with Dunnett’s post-test comparing all evolved lineages to the ancestor. Error bars represent standard deviation. * p<0.05, ** p<0.01, *** p<0.001.

Another *P. aeruginosa* phenotype assessed was biofilm formation. Based on the design of our experimental evolution, we hypothesized that the evolved lineages would be superior biofilm formers compared to their ancestors because our experimental evolution design strongly selected for cells that were adhered to the tissue culture surface. We assessed biofilm formation with crystal violet assays which provide a broad quantitative means to assess the volume of biofilm biomass produced.

Surprisingly, only two strains, PA B80398 and PA3, exhibited significantly different biofilm densities compared to their ancestor strains (Fig. 3). In both instances, the evolved lineages exhibited significantly reduced biofilm biomass compared to their ancestors, though the overall degree of loss is quite small (5-25%). This result is consistent when considering each evolved lineage separately as all lineages showed reduced biofilm biomass (Fig. S3) which suggests that this change was not driven by data from a single lineage but likely represents a true shift in the phenotype for these strains. All the remaining evolved lineages tested were not significantly different from their ancestor with respect to biofilm formation (Fig. 3 and S3).

**Figure 3.**
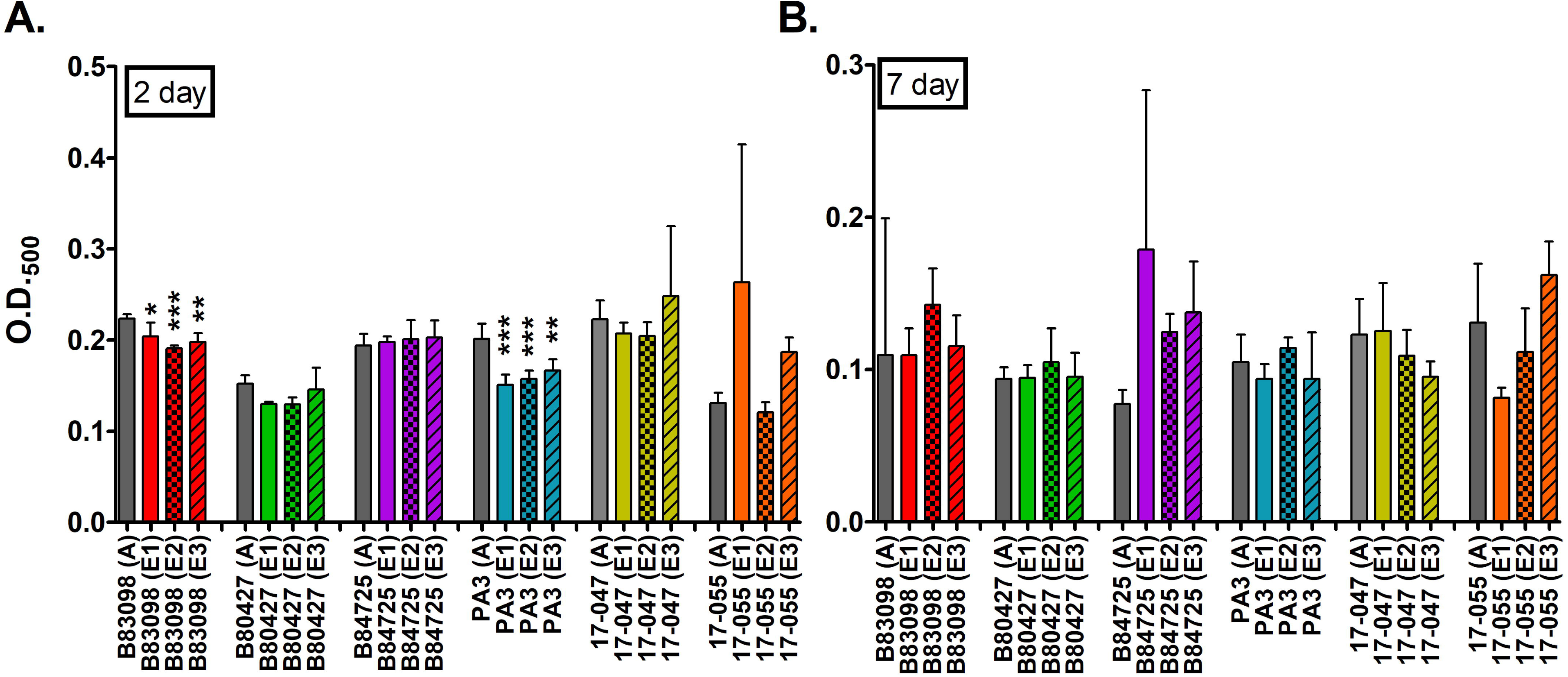
Biofilm formation of ancestral and evolved isolates in 1% HL5. Panels show crystal violet retention after (A) 2 days or (B) 7 days of incubation. Note the different y-axis scales. Grey bars represent ancestors and colored bars represent evolved isolates. Each bar represents the average of three biological replicates for each isolate. An (A) in the x-axis indicates the ancestral isolate, and (E) represents evolved isolates. Data analyzed with t-tests between the ancestor and evolved lineages. Error bars represent standard deviation. * p<0.05, ** p<0.01, *** p<0.001.

### Individual molecular phenotypes

Pyocyanin is a redox-active toxin secreted by *P. aeruginosa.* It is toxic to eukaryotic cells and provides *P. aeruginosa* protection against predators, but it also serves as a secondary metabolite for *P. aeruginosa* and is associated with promoting survival in low-oxygen and low-nutrient environments (48–50). We hypothesized that the evolved lineages would produce significantly more pyocyanin than their ancestors due to the low-nutrient conditions present in our experimental evolution system. Our hypothesis was supported for three of the strains: PA B80427, SRP 17-047, and SRP 17-055, of which there was a small but significant increase in pyocyanin production in PA B80427 and SRP 17-047, and a large increase in SRP 17-055 (Fig. 4). Two strains, PA B80398 and PA B84725, did not exhibit significant differences in pyocyanin production compared to their ancestors; however, two of the individual evolved lineages of PA B80398 did produce more pyocyanin than their ancestor (Fig. S4). Interestingly, all three evolved lineages of PA3 produced significantly less pyocyanin compared to their ancestor suggesting that pyocyanin production was not selected for in this strain. The reasons for this remain unclear. From this data, it suggests that for most of our *P. aeruginosa* strains pyocyanin production was selected for in our experimental evolution system and increased pyocyanin increased the fitness of the organisms in these low-nutrient environments.

**Figure 4.**
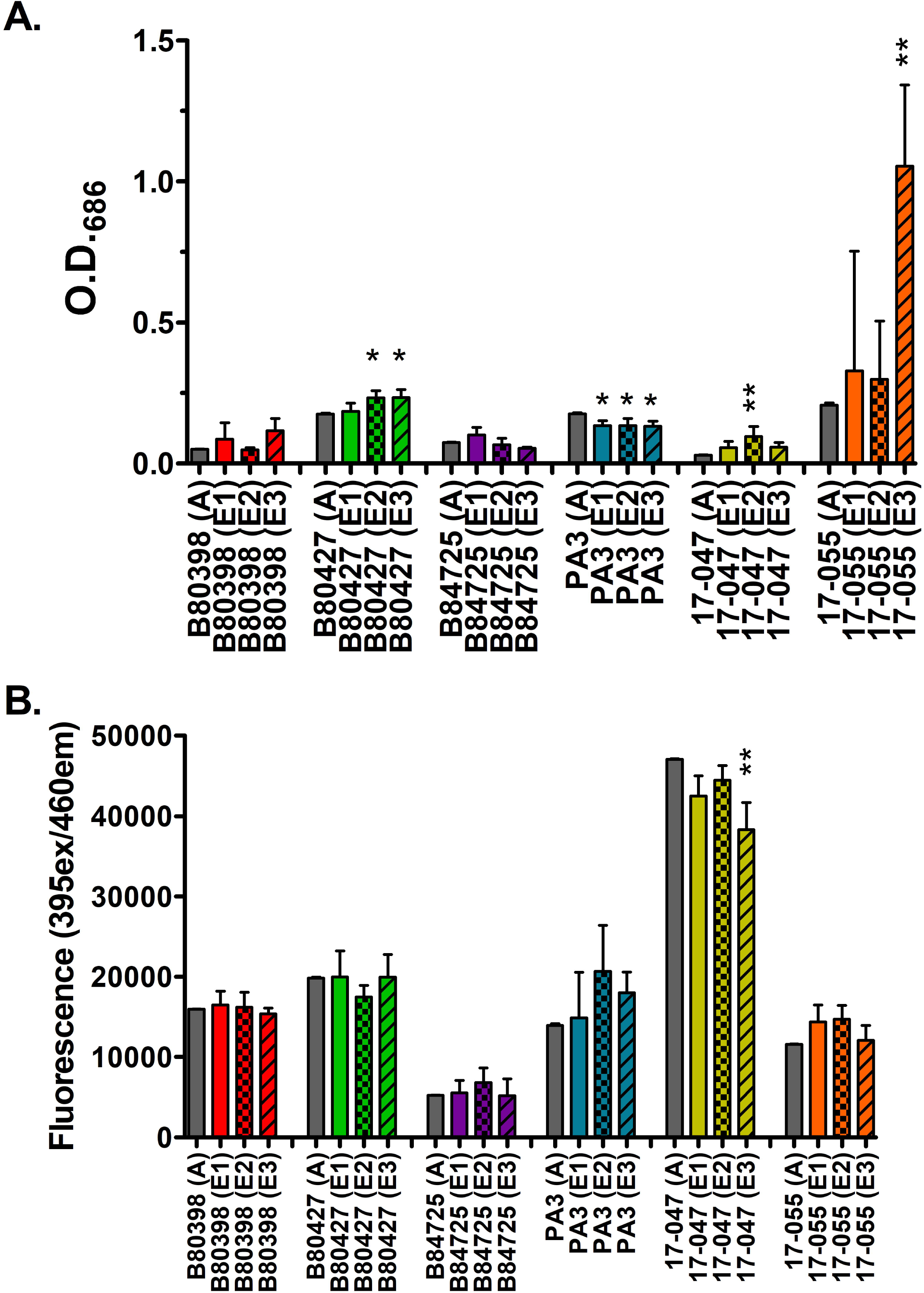
Pigment production in ancestral and evolved *P. aeruginosa* isolates. Panels show (A) pyocyanin or (B) pyoverdine production from 12-week-old evolved lineages compared to ancestors. Grey bars represent ancestors and colored bars represent the evolved lineages. Each bar represents the average of three biological replicates for each isolate. An (A) in the x-axis indicates the ancestral isolate, and (E) represents evolved isolates. Data analyzed with t-tests between the ancestor and evolved lineages. Error bars represent standard deviation. * p<0.05, ** p<0.01.

Pyoverdine is a siderophore secreted by *P. aeruginosa* which functions to sequester iron from other cells as well as the environment (51). The 1% HL5 used for the experimental evolution is undoubtedly low in most macronutrients. However, it likely does contain some iron as it contains yeast extract and protease peptone but whether the concentration was sufficient for *P. aeruginosa* growth is unknown. Therefore, we could not make a hypothesis on whether pyoverdine production would change over time in the evolved isolates. Pyoverdine is highly implicated in *P. aeruginosa* pathogenicity (52) so pyoverdine production was assessed using spectrophotometry in the ancestral and evolved lineages.

Pyoverdine remained unchanged in five of the six strains tested (Fig. 5). Only one of the evolved lineages of the environmental isolate, SRP 17-047 produced significantly less pyoverdine than its ancestor, while the other two lineages of 17-047 were not significantly different (Fig. S5). This suggests that pyoverdine was likely neither selected for nor against under the conditions in our experimental evolution system.

**Figure 5.**
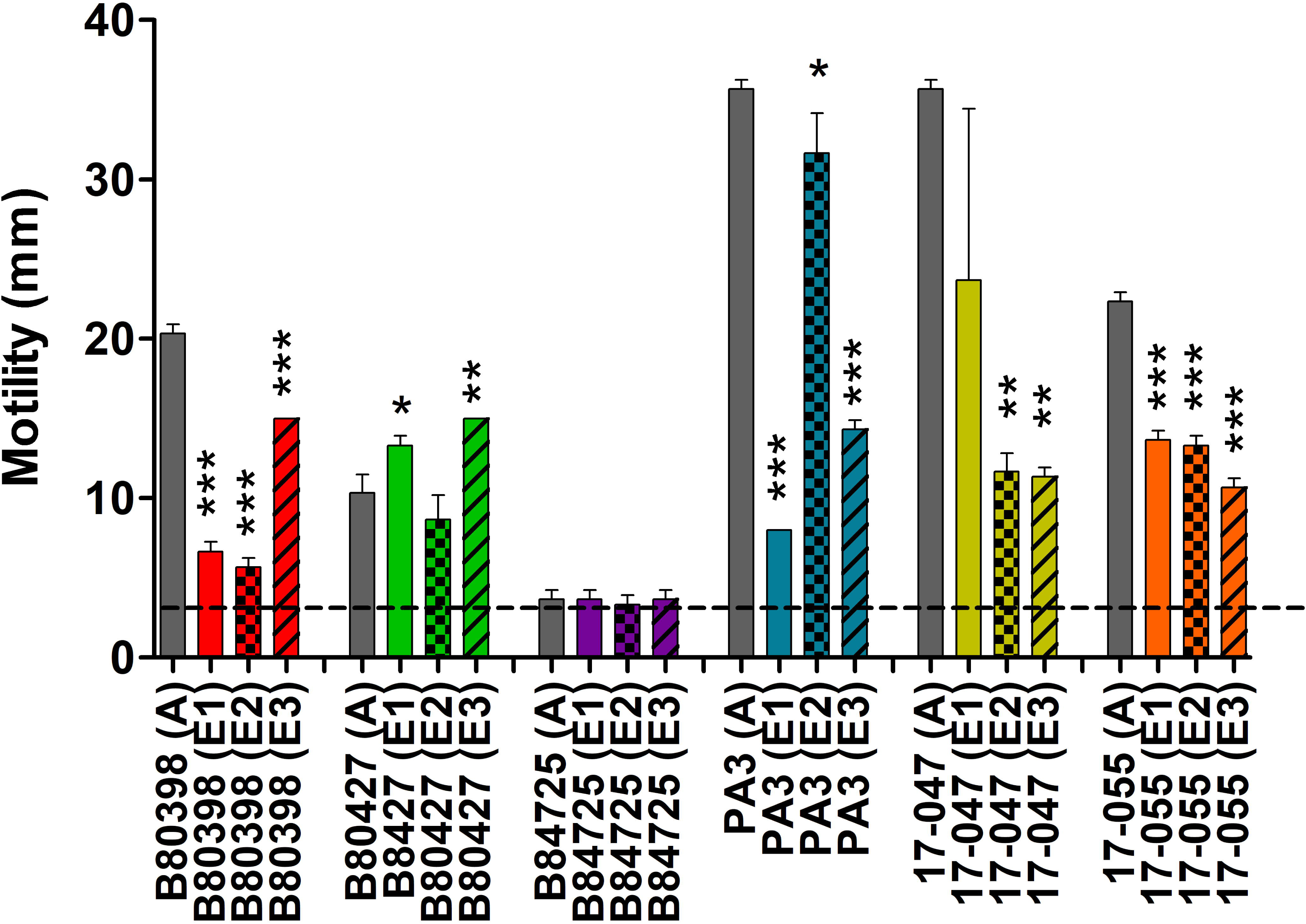
Swimming motility of ancestral and 12-week evolved isolates in 0.3% LB agar. Grey bars represent ancestors and colored bars represent evolved isolates. Each bar represents the average of three biological replicates for each isolate. The dashed line indicates the diameter of the stab mark. An (A) in the x-axis indicates the ancestral isolate, and (E) represents evolved isolates. Data analyzed with t-tests between the ancestor and evolved lineages. Error bars represent standard deviation. * p<0.05, ** p<0.01, *** p<0.001.

The last molecular phenotype tested was swimming motility. We hypothesized that there would be a reduction in motility in our evolved lineages as our experimental evolution design selected against planktonic cells. The swimming abilities of the ancestral isolates were variable from completely non-motile to highly motile (Fig. 5). There was a significant reduction in swimming motility observed for almost all the evolved lineages compared to their ancestors for B80398, PA3, 17-047, and 17-055. The other two strains, B80427 and B84725, showed very small changes in motility compared to their ancestors; however, the ancestors for these two strains were either amotile (B84725), or only weakly motile (B80427), under these conditions. Surprisingly, two of the evolved lineages, E1 and E3 of B80427, displayed small but significantly greater motility than the ancestor B80427 strain. There was no correlation between biofilm biomass and motility for all strains aggregated (Spearman r, *p*=0.5720) or for any strain individually.

We tested the earlier isolates of the four strains that showed the greatest reduction in motility over the course of the experiment, B80398, PA3, 17-047, and 17-055, to determine when motility was lost by selecting isolates from earlier time points in the experimental evolution. Isolates from the second week of the experimental evolution were tested first, then isolates from additional weeks were tested as needed. We found that motility was lost at week 2 for 17-047, week 3 for 17-055, week 5 for PA3, and week 7 for B80398 (Fig. S5). Together these data suggest that for strains that were highly motile at the start of the experiment, motility was selected against and loss of 49-90% of motility between 2 and 7 weeks of culture in low-nutrient medium. The degree and timing of the loss varied for each motile strain.

### Host-associated phenotypes over the course of evolution in a low nutrient environment

We also sought to determine if long-term adaptation to a low-nutrient environment would result in an increased or decreased ability to survive and/or kill a phagocytic predator as this could be a strong indicator of how *P. aeruginosa* virulence can increase in low-nutrient environments. To test this, we co-cultured the ancestral isolates and one evolved lineage of each strain with the phagocytic amoeba species *A. castellanii* over time to examine the survival of both organisms*. A. castellanii,* an avid predator of *P. aeruginosa,* is a model organism for studying phagocytosis and is often used as a proxy to study mammalian macrophages engulfment (53–56). *A. castellanii* exists in two forms, trophozoites (the actively growing and metabolizing form) and cysts (dormant).

We examined trophozoite and *P. aeruginosa* survival for individual ancestral and evolved strains. After 16 days of co-culture, *A. castellanii* trophozoite survival was significantly reduced when co-cultured with the evolved lineage of the clinical isolates B80398 and PA3 when compared with the ancestral isolate of these strains (Fig. 6A). The remaining strains showed neither a statistical increase nor decrease in the number of trophozoites found, perhaps due to the large variability in the individual lineages for the most part. Trophozoite survival tended to be reduced over time when in co-culture with the evolved lineage compared to the ancestral isolates for most strains but this rarely reached statistical significance for most lineages/strains (Fig. S6). The co-culture dynamics of the clinical strains PA3 and B80398 over time were particularly interesting. Trophozoite levels in presence of the ancestral strains increased dramatically over the 16 days of observation but trophozoite levels remained very low when co-cultured with the evolved isolate of this strain (Fig. S6).

**Figure 6.**
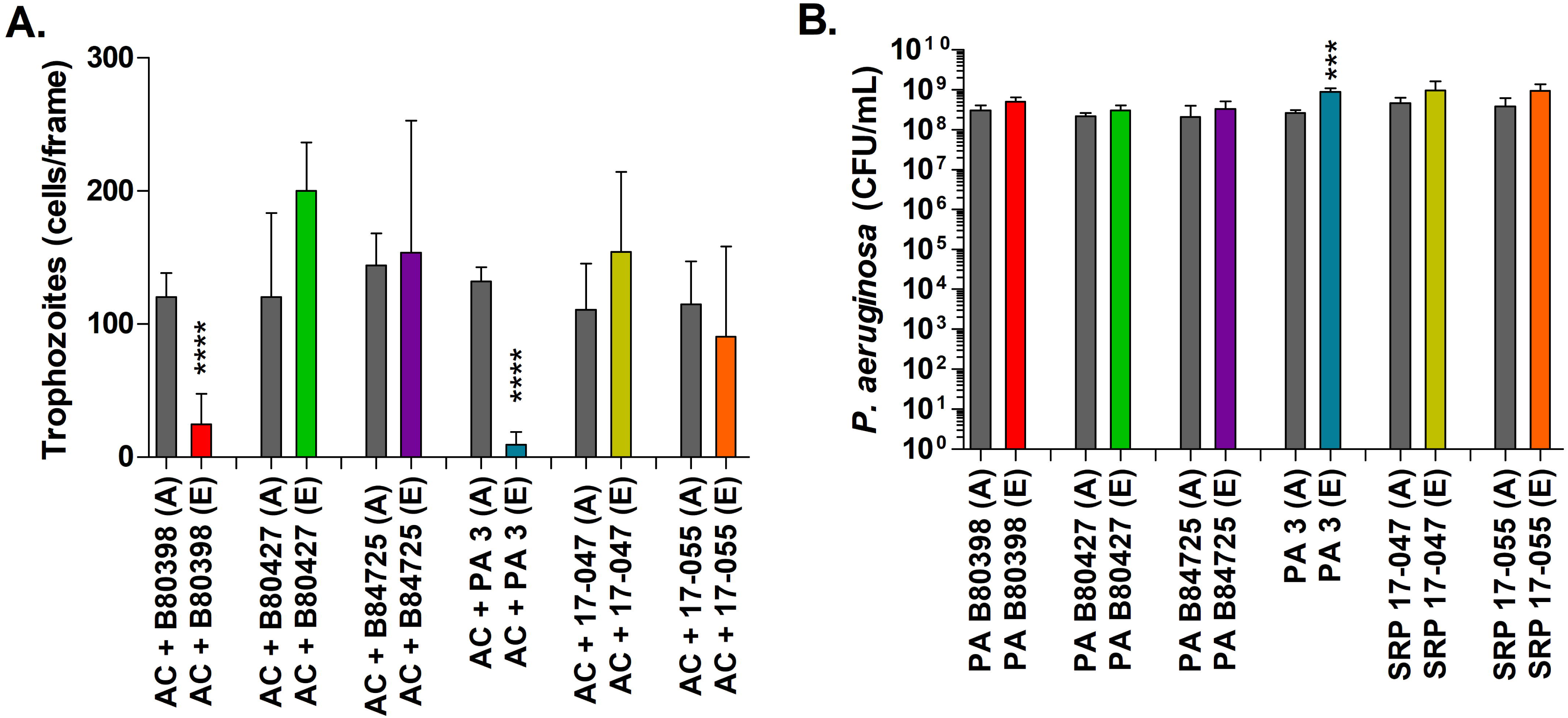
*A. castellanii* and *P. aeruginosa* survival after 16 days of co-culture. (Panel A) Bars represents the average number of A. castellanii trophozoites counts from microscopy images. (Panel B) *P. aeruginosa* viable cell counts from ancestral or a single evolved lineage in each co-culture condition (line 3 for strain SRP 17-047 and line 2 for strains B80398, B80427, B84725, PA3, and SRP 17-055) are shown. Grey bars represent ancestors (A) and colored bars represent the evolved lineage (E). Error bars represent standard deviation (n=3). Data analyzed with t-tests between ancestor and evolved isolates. *** p<0.0005, **** p<0.0001.

In comparison, all the *P. aeruginosa* isolates tested were able to survive in co-culture with *A. castellanii* for at least 16 days (Fig. 6B). There were no significant differences between the numbers of surviving evolved cells compared to ancestors at 16 days except for the clinical isolate, PA3, for which the number of surviving cells at 16 days was significantly greater for the evolved lineage than the ancestor. Therefore, ancestral and evolved lineages of *P. aeruginosa* were able to persist in the presence of amoebae. *P. aeruginosa* cells were only enumerated on day 16 as we were able to confirm the presence of live *P. aeruginosa* cells in the wells using bright-field microscopy at each of the other time points.

### Genomic changes in the evolved lineages compared to ancestors

To better understand the genetic mechanisms underlying the observed changes in phenotype and to better identify other potential changes, the genomes of the ancestor and one of the evolved lineages from four of the strains, B83098, B80427, 17-047, and 17-055, were sequenced and compared. The genomes sizes ranged from 6.60 Mbp to 6.99 Mbp (average 6.81 Mbp) and the number of predicted genes was between 6149 and 6456 (average 6,318) (Table 2). The number of SNPs in coding regions between ancestor and evolved isolates varied between 212 (17–055) and 312 (B80427).

**Table 2.**
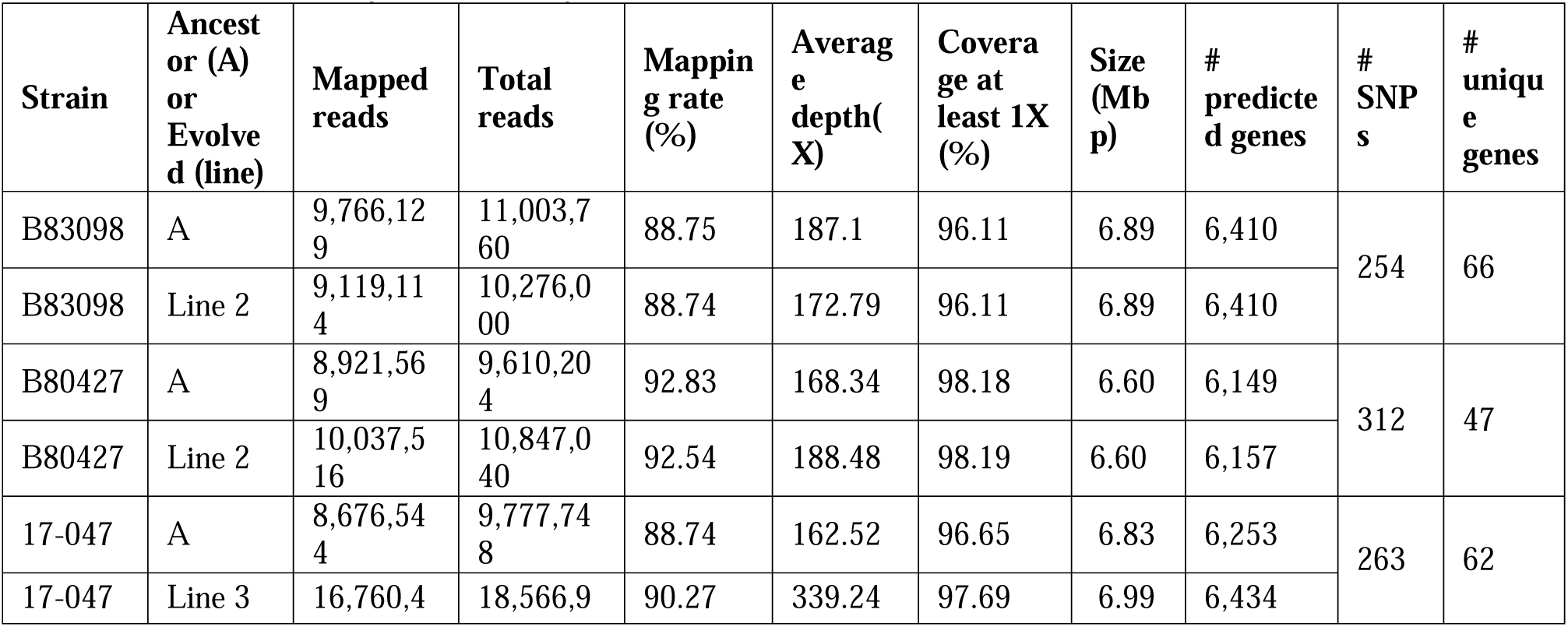
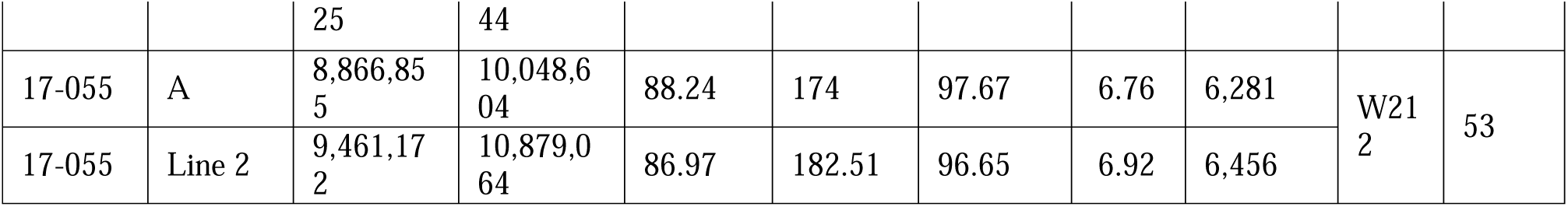
Genome analysis summary.

We identified coding genes that contained SNPs between ancestral and evolved lineages (Fig. 7). There were 6 genes or homologous gene pairs that were mutated in all four evolved lineages. They encoded: phenazine biosynthesis proteins PhzA2/PhzA1; Type VI secretion system tip proteins VgrG/VgrG1; a short chain dehydrogenase; outer membrane protein OprM; glutamate 5-kinase (involved in the urea cycle); and 3-carboxy-cis,cis-muconate cycloisomerase (part of the protocatechuate pathway that is a central catabolic route for aromatic compounds). A number of other genes were mutated in 3 of the four strains. Notably, genes that encode putative virulence factors like VirS [a transcriptional regulator of virulence proteins] and TpsA/B proteins [involved in two partner secretion of virulence proteins (57)], genes that encode histidine biosynthesis proteins. There were no genes containing SNPs that were only shared between B80398 and the two environmental strains, 17-047 and 17-055.

**Figure 7.**
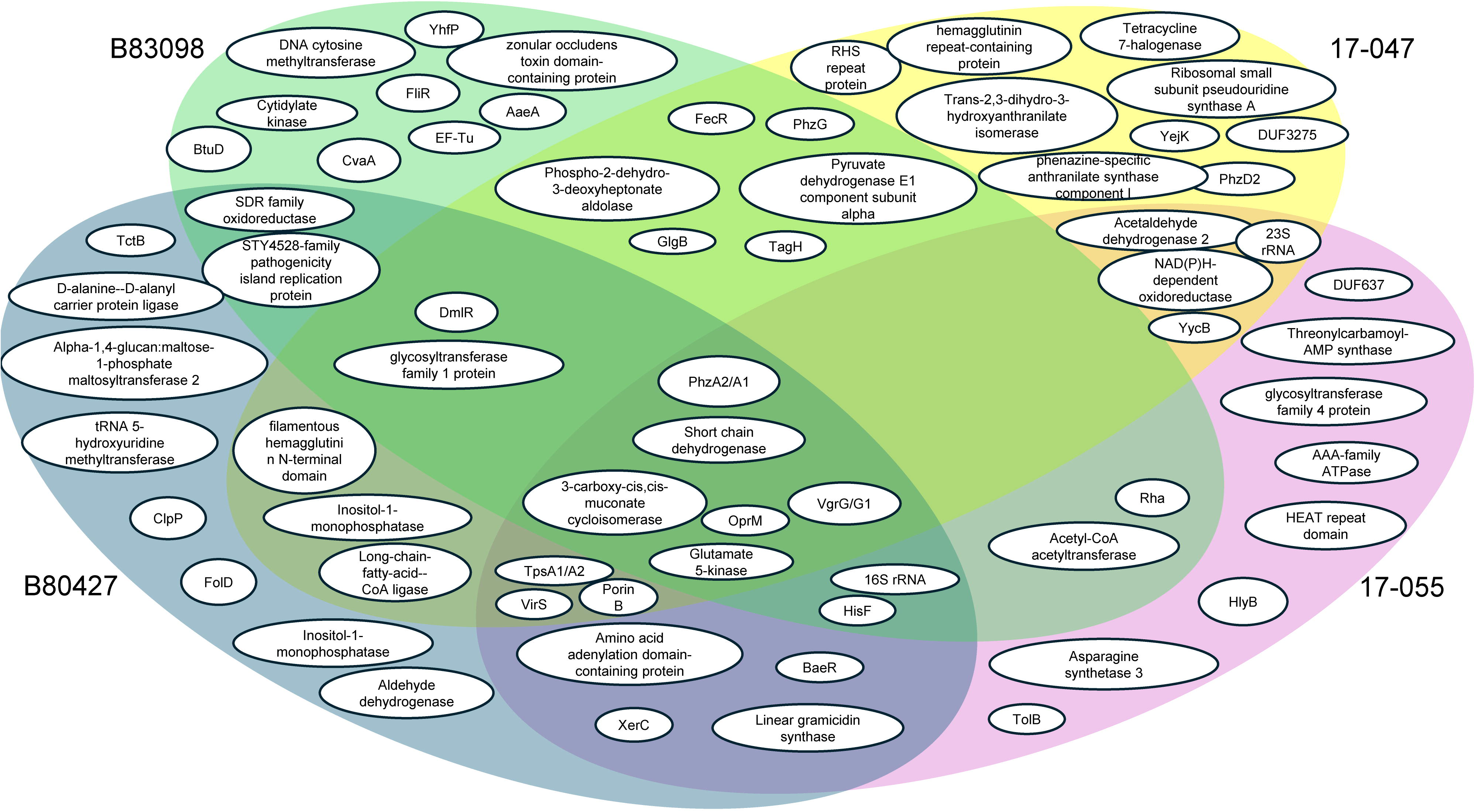
Venn diagram of single nucleotide polymorphisms between ancestors and their respective evolved lineages after the 12-week experimental evolution. Genes with at least two SNPs in at least one of the evolved lineages are represented.

Additionally, SNPs that were shared between clinical strains or environmental strains were also identified. The clinical isolates B80398 and B80427 had only two unique coding regions with SNPs – a SDR oxidoreductase gene and a STY4528-family pathogenicity island replication protein. The environmental strains showed SNPs in four different genes, encoding acetylaldehyde 2, a NAD(P)H-dependent oxidoreductase, the 23S rRNA, and a gene that was annotated as YycB, a uncharacterized transporter from *Bacillus subtilis*.

In the molecular phenotype analyses, motility showed the greatest level of changes between ancestral and evolved lineages. We assessed the genomic DNA results and found that only one strain, B80398, had SNPs in a known flagella-associated protein. FliR is a core part of the FliP/FliQ/FliR complex that forms a proton motive force-dependent channel that exports flagellar proteins across the cytoplasmic membrane (58) and has also been implicated in the secretion of virulence factors (59) and in adhesion and biofilm formation (60) in other organisms. DmlR, a transcription factor implicated in carbohydrate metabolism and flagellar motility regulation contained SNPs in three of the four strains, B80398, B80427, and 17-047 (61) though this association is only hypothetical. Evolved lineages of these strains may also have SNPs in intergenic, regulatory regions or in other types of regulatory proteins that could explain the changes in motility observed.

## Discussion

The adaptations that occur in response to long periods without sufficient food sources could impact how bacteria grow (both in terms of how fast and how to divide their limited resources), sense and respond to their environment, their lifestyle, or how they defend themselves against predation. Culturing *P. aeruginosa* for 3 months under low-nutrient conditions resulted in alterations to multiple phenotypes associated with fitness and/or virulence. Some of these observed changes in phenotype were shared amongst most of the evolved lineages, regardless of ancestral strain. These changes in phenotypes include faster generation time, reduced cell size, increased pyocyanin production, and decreased motility. Since these phenotypes were observed in most lineages, it could be inferred that these phenotypic alterations lead to increased fitness in low nutrients and thus are likely to occur in these types of environments. Two other phenotypes tested were biofilm formation and pyoverdine production, and we observed variable changes between strains and within replicate evolved lineages and within each strain. This indicates that these phenotypes were likely not as strongly selected for in our experimental evolution system (62).

The environments where these particular *P. aeruginosa* strains were isolated likely had higher levels of nutrients (i.e. the clinical isolates) or fluctuating levels of nutrients (household drains). Therefore, the ancestral strains used in our experiments were likely not well adapted to constant low nutrient conditions. Evolved lineages of three of the six strains tested exhibited significantly faster growth compared to their ancestors (at either 22°C, 37°C, or both), indicating that the evolved lineages became better adapted to the environment in which they were evolved (Fig. 1). These results are consistent with results observed in other experimental evolution systems such as those observed in the Long-Term Evolution Experiment (LTEE) of *Escherichia coli* where the doubling times of the evolved lineages reduced from 55 minutes to 23 minutes in the minimal nutrient medium used in the experiment after ∼50,000 generations (62). In contrast, a few of the evolved isolates (e.g. B80398 evolved line 2) exhibited significantly increased generation times compared to their ancestor (Fig. 1). Surprisingly, we did not observe any direct correlation between cell size and generation time suggesting that cell size does not play a factor in doubling time for *P. aeruginosa* as it does for *E. coli* (46), at least not over the short-term evolution experiment described here.

Our experimental evolution system theoretically was expected to have strongly selected for increased biofilm formation. Surprisingly, evolved lineages in two of the six strains produced significantly less biofilm biomass than their ancestors while the evolved lineages of the remaining four strains were not significantly different from their ancestors (Fig. 3). One potential explanation for this could be the low-nutrient culture medium used in the evolution experiment. Biofilms require the production and secretion of polysaccharides and extracellular proteins to form the biofilm matrix (63–65). These matrix components are undoubtedly metabolically costly to produce as the responsibility for exopolysaccharide matrix generation has been shown to be a shared responsibility within a biofilm in other species of bacteria such as *Bacillus subtilis* (66). Thus, it is possible that the cells either acquired mutations that blocked the biosynthesis or export of some of the necessary biofilm components – perhaps choosing to use the limited carbon for growth rather than matrix production. In addition, the evolved cells could have used other methods of attachment to the plate surface, such as pili or fimbriae, to avoid the expense of secreting polysaccharides though no SNPs were found in the coding regions of known pilin or fimbriae genes. Alternatives could include proteins like FliR, which has been shown to be involved in biofilm and adhesion (60) or be due to SNPs in transcriptional factors. Thus, there are several alternatives that could explain how *P. aeruginosa* could have attached to the surface but not initiated polysaccharide secretion. The lack of increase in biofilm formation in low-nutrient environments is similar to those observed in *P. aeruginosa* strains co-cultured long-term with the amoebal predator, *A. castellanii,* in a minimal nutrient medium (67). In this latter study, biofilm biomass was also found to decrease in *P. aeruginosa* strains cultured in monoculture as well as in co-culture with *A. castellanii* in minimal medium for 42 days.

We expected that flagellar motility would be selected against in our experimental evolution because planktonic cells were removed during every passage. A reduction in motility was observed in the evolved lineages of strains where the ancestors exhibited high motility (Fig. 5). This result is similar to results observed in *P. aeruginosa* PAO1 cultured in artificial CF medium for 120 days under aerated conditions both in monoculture and in the presence of phage competitors (68). In our study, the evolved lineages of two strains, PA B80427 and PA B84725, did not exhibit a reduction in motility, perhaps because the ancestors were poorly motile/amotile and possibly do not have sufficient loci for evolution to act upon. We did not observe a significant correlation, either negatively or positively, between biofilm formation and motility as others have previously (69, 70). The reasons for this remain unclear though there have been a few studies that also did not find correlations between these phenotypes in *P. aeruginosa*, particularly when *P. aeruginosa* forms aggregates (71, 72), or suggest that the flagellar dependence for biofilm formation may depend on the growth medium (73).

We also had hypothesized that pyocyanin production would be increased in the evolved populations compared to their ancestors because outside of its toxic effects on host and other prokaryotic cells, pyocyanin serves as a secondary metabolite, stabilizing the redox state of *P. aeruginosa* and can act as a final electron acceptor in the electron transport chain (38). Increased expression/production of pyocyanin should therefore increase metabolic fitness in low nutrient environments. Pyocyanin has also been shown to play a role in pyruvate excretion and metabolism under anoxic conditions which are common in the lung environment and in biofilms (48–50). *P. aeruginosa* has been shown to be able to ferment excreted pyruvate to survive in anoxic/low-nutrient conditions for at least 18 days (49). Our pyocyanin hypothesis was supported for half of the evolved lineages when three of the six strains showed a significant increase in pyocyanin production compared to their ancestors. Because our cultures were static, it could be that increases in pyocyanin were observed for those strains that needed additional electron acceptors for their particular electron transport chain while other strains did not.

In contrast, production of pyoverdine, a siderophore, was not significantly altered in almost all evolved lineages compared to their ancestors. This result indicates that pyoverdine was not selected for or against in our experimental evolution design. This result is in opposition to results observed in previous studies which found that pyoverdine production decreased over the course of 42 days in a minimal nutrient medium (67). It is possible that the 1% HL5 used in our study contained more freely available iron and thus was not selected for or against in our experimental evolution system compared to the supplemented M9 medium used in the previous study.

In addition to identifying phenotypic changes that occurred in our evolved isolates, we also sought to determine if adaptations to a low-nutrient environment could have implications on virulence against host organisms. *A. castellanii* a bacterivorous amoebae species was identified as a model organism to study phagocytosis because of its commonalities with phagocytic human immune cells, such as macrophages (53–56). In our study, trophozoite survival was sometimes decreased when co-cultured with evolved isolates compared to ancestors, particularly strains B83098 and PA3, but never significantly increased when co-cultured with the ancestral isolates (Figs. 6 & S6). Additionally, all the evolved lineages of *P. aeruginosa* were just as capable of surviving predation from the amoebae predator as their ancestors were (Fig. 6B). One potential reason that might underlie why *A. castellanii* trophozoites tended to survive better with the ancestors than with the evolved lineages over time could be due to the reduced motility observed in our evolved lineages. *Campylobacter jejuni, Pseudomonas fluorescens,* and *Proteus mirabilis* flagellin are recognized by *A. castellanii* to initiate phagocytosis (74, 75). Thus, if there are fewer/less flagella expressed in the evolved isolates, there would be less predation by *A. castellanii* resulting in starvation of the amoebae predator when in co-culture with the evolved lineages.

The genomes of the evolved and ancestral isolates showed many differences in the presence of SNPs. Whether or not these SNPs actually relate to changes in the amino acid structure of a protein or whether they influence the expression of these genes remains to be seen. However, understanding the precise mutations that occurred can shed light on the genes where natural selection may be occurring. We also note that individual isolates of evolved lines were chosen for genome sequencing, and this may have limited the resolution of the SNPs that occurred across the population. Previous studies have shown that long term starvation, nutrient exhaustion, or feast-and-famine lifestyles can lead to increased genomic heterogeneity in bacterial populations rather than communities dominated by one phenotype produced from beneficial SNP(s) (76–78). Further, we observed evidence of mutational parallelism among the strains after 12 weeks as was observed in (78), despite the feast and famine cycles somewhat mirrored in our study were much shorter than the 100 day cycles used in that study. For example, strains PA B80398 and PA 80427 showed mutations in the gene encoding the STY45280family pathogenicity island replication protein and

In summary, a variety of *P. aeruginosa* phenotypes were observed to change over the course of long periods with limited resources, but there was a great deal of variability in how *P. aeruginosa* chose to invest its resources based on individual lineage data. This may not be surprising, given the genomic plasticity within *P. aeruginosa* as a species. These results are significant as they contribute to fundamental scientific knowledge into how *P. aeruginosa* might evolve in other low nutrient environments (such as sink drains and faucets, hospital surfaces, or freshwaters). This work could also yield insights into expected changes in *P. aeruginosa* over time in low-nutrient environments and this could have implications for virulence properties or treatment efficacy for human infections. It could help us to understand the mechanisms of adaptation in such non-host environments (as in static cultures) and optimize sanitation protocols, especially in healthcare settings, and/or designing surfaces or materials that inhibit bacterial attachment and persistence. Lastly, this work could set the stage for future evolution experiments in which *P. aeruginosa* could be evolved with predators, such as macrophage or amoeba, to examine the effect of nutrient concentration on host-pathogen interactions.

## Supplemental figure legends

**Figure S1.** Growth curves of the evolved lineages and their ancestors over time in 1% HL5 at room temperature (panels A-F) or 37°C (panels G-L). Lines represent the mean of 3 biological replicates. Error bars are not shown for image clarity. Data analyzed with repeat measures one-way ANOVA with Dunnett’s post-test comparing evolved lineages to their ancestor. * p<0.05, ** p<0.01, *** p<0.001.

**Figure S2.** Cell size of individual evolved lineages compared to their ancestor are shown with each cell measure represented by a dot. Cells grown at room temperature (panels A-F) or 37° (panels G-L) to exponential phase were measured. Panels M and N show aggregated data for ancestral (A) and evolved (E) isolates. Horizontal bars for each strain represent the median cell size of 100-600+ cells and error bars represent the interquartile range. Data were analyzed with one-way ANOVA with Dunnett’s post-test comparing each evolved line to its ancestor. * p<0.05, ** p<0.01, *** p<0.001, **** p<0.0001.

**Figure S3.** Biofilm biomass of individual evolved lineages compared to their ancestor after 3-month experimental evolution. Data for evolved lineages was aggregated to assess global patterns of evolution. Error bars represent standard deviation from four independent colonies from each lineage. Data analyzed with one-way ANOVA with Dunnett’s posttest comparing each evolved line to its ancestor. *** p<0.001, **** p<0.0001.

**Figure S4.** Aggregate data for evolved *P. aeruginosa* lineages for pyocyanin (panel A) or pyoverdine (panel B) production of individual isolates compared to their ancestor after the 12-week experimental evolution. Error bars represent standard deviation from three independent colonies from each lineage. Data analyzed with one-way ANOVA with Dunnett’s post-test comparing each evolved line to its ancestor. * p<0.05, ** p<0.01.

**Figure S5.** Swimming motility of individual isolates compared to their ancestor for isolates frozen at various points during the experimental evolution. Week 2 isolates were first tested, then week 3, 5, and/or 7 as needed until the loss of motility was observed. Error bars represent standard deviation from three independent colonies from each lineage. 35 mm was the largest measurable size in this analysis. N.D. indicates non-detectable growth past the stab line. Data analyzed with two-way ANOVA with Bonferroni post-test comparing each evolved line to its ancestor across time points. Error bars indicate standard deviation. * p<0.05, ** p<0.01, *** p<0.001, **** p<0.0001.

**Figure S6.** Competition assay between the ancestor and evolved lineages (line 3 for strain SRP 17-047 and line 2 for strains B80398, B80427, B84725, PA3, and SRP 17-055) of *P. aeruginosa* against *A. castellanii* at all timepoints tested. Grey bars with or colored bars represent the number of trophozoites observed when co-cultured with each ancestor (A in the x-axis) or evolved lineage (E in the x-axis) over the course of 16 days (D) Panels represent each strain with (A) B80398, (B) B80427, (C) B84725, (D) PA3, (E) 17-047, and (F) 17-055 measured at 0-, 3-, 6-, 8-, 11-, 13-, and 16 days of co-culture. Error bars represent standard deviation from three independent colonies from each lineage. Data analyzed with t-tests between ancestor and evolved lineage at each timepoint. * p<0.05, ** p<0.01, *** p<0.001, **** p<0.0001.

## References

1. Zhu M, Dai X. 2023. Stringent response ensures the timely adaptation of bacterial growth to nutrient downshift. Nat Commun 14:467.

2. Jain V, Kumar M, Chatterji D. 2006. ppGpp: stringent response and survival. J Microbiol 44:1–10.

3. Wang X, Wang J, Liu SY, Guo JS, Fang F, Chen YP, Yan P. 2023. Mechanisms of survival mediated by the stringent response in *Pseudomonas aeruginosa* under environmental stress in drinking water systems: Nitrogen deficiency and bacterial competition. J Hazard Mater 448:130941.

4. Gray DA, Dugar G, Gamba P, Strahl H, Jonker MJ, Hamoen LW. 2019. Extreme slow growth as alternative strategy to survive deep starvation in bacteria. Nat Commun 10:890.

5. Errington J. 1993. *Bacillus subtilis* sporulation: regulation of gene expression and control of morphogenesis. Microbiol Rev 57:1–33.

6. Huberts DH, Niebel B, Heinemann M. 2012. A flux-sensing mechanism could regulate the switch between respiration and fermentation. FEMS Yeast Res 12:118–28.

7. van Hoek MJ, Merks RM. 2012. Redox balance is key to explaining full vs. partial switching to low-yield metabolism. BMC Syst Biol 6:22.

8. Santajit S, Indrawattana N. 2016. Mechanisms of Antimicrobial Resistance in ESKAPE Pathogens. Biomed Res Int 2016:2475067.

9. Hancock RE, Speert DP. 2000. Antibiotic resistance in *Pseudomonas aeruginosa*: mechanisms and impact on treatment. Drug Resist Updat 3:247–255.

10. Fernandez-Billon M, Llambias-Cabot AE, Jordana-Lluch E, Oliver A, Macia MD. 2023. Mechanisms of antibiotic resistance in *Pseudomonas aeruginosa* biofilms. Biofilm 5:100129.

11. Botelho J, Tuffers L, Fuss J, Buchholz F, Utpatel C, Klockgether J, Niemann S, Tummler B, Schulenburg H. 2023. Phylogroup-specific variation shapes the clustering of antimicrobial resistance genes and defence systems across regions of genome plasticity in *Pseudomonas aeruginosa*. EBioMedicine 90:104532.

12. Mendes OR. 2023. Chapter 48 - The challenge of pulmonary *Pseudomonas aeruginosa* infection: How to bridge research and clinical pathology, p 591–608. In Bagchi D, Das A, Downs BW (ed), Viral, Parasitic, Bacterial, and Fungal Infections 10.1016/B978-0-323-85730-7.00019-9. Academic Press.

13. Walker TS, Bais HP, Deziel E, Schweizer HP, Rahme LG, Fall R, Vivanco JM. 2004. *Pseudomonas aeruginosa*-plant root interactions. Pathogenicity, biofilm formation, and root exudation. Plant Physiol 134:320–31.

14. Thi MTT, Wibowo D, Rehm BHA. 2020. *Pseudomonas aeruginosa* Biofilms. Int J Mol Sci 21.

15. Britigan BE, Roeder TL, Rasmussen GT, Shasby DM, McCormick ML, Cox CD. 1992. Interaction of the *Pseudomonas aeruginosa* secretory products pyocyanin and pyochelin generates hydroxyl radical and causes synergistic damage to endothelial cells. Implications for *Pseudomonas*-associated tissue injury. J Clin Invest 90:2187–96.

16. Hauser AR. 2009. The type III secretion system of *Pseudomonas aeruginosa*: infection by injection. Nat Rev Microbiol 7:654–65.

17. Hogardt M, Heesemann J. 2010. Adaptation of *Pseudomonas aeruginosa* during persistence in the cystic fibrosis lung. Int J Med Microbiol 300:557–62.

18. Frimmersdorf E, Horatzek S, Pelnikevich A, Wiehlmann L, Schomburg D. 2010. How *Pseudomonas aeruginosa* adapts to various environments: a metabolomic approach. Environ Microbiol 12:1734–47.

19. Ebert D. 1998. Experimental evolution of parasites. Science 282:1432–5.

20. Wong A, Rodrigue N, Kassen R. 2012. Genomics of adaptation during experimental evolution of the opportunistic pathogen *Pseudomonas aeruginosa*. PLoS Genet 8:e1002928.

21. Bakkal S, Robinson SM, Ordonez CL, Waltz DA, Riley MA. 2010. Role of bacteriocins in mediating interactions of bacterial isolates taken from cystic fibrosis patients. Microbiology (Reading) 156:2058–2067.

22. Merritt JH, Kadouri DE, O’Toole GA. 2005. Growing and analyzing static biofilms. Curr Protoc Microbiol Chapter 1:Unit 1B 1.

23. Gajdacs M, Barath Z, Karpati K, Szabo D, Usai D, Zanetti S, Donadu MG. 2021. No Correlation between Biofilm Formation, Virulence Factors, and Antibiotic Resistance in *Pseudomonas aeruginosa*: Results from a Laboratory-Based In Vitro Study. Antibiotics (Basel) 10.

24. S A. 2010. FastQC: A Quality Control Tool for High Throughput Sequence Data. http://www.bioinformatics.babraham.ac.uk/projects/fastqc/. Accessed

25. Martin M. 2011. Cutadapt removes adapter sequences from high-throughput sequencing reads. 2011 17:3.

26. Bolger AM, Lohse M, Usadel B. 2014. Trimmomatic: a flexible trimmer for Illumina sequence data. Bioinformatics 30:2114–20.

27. Li H, Durbin R. 2009. Fast and accurate short read alignment with Burrows-Wheeler transform. Bioinformatics 25:1754–60.

28. Danecek P, Bonfield JK, Liddle J, Marshall J, Ohan V, Pollard MO, Whitwham A, Keane T, McCarthy SA, Davies RM, Li H. 2021. Twelve years of SAMtools and BCFtools. Gigascience 10.

29. McKenna A, Hanna M, Banks E, Sivachenko A, Cibulskis K, Kernytsky A, Garimella K, Altshuler D, Gabriel S, Daly M, DePristo MA. 2010. The Genome Analysis Toolkit: a MapReduce framework for analyzing next-generation DNA sequencing data. Genome Res 20:1297–303.

30. DePristo MA, Banks E, Poplin R, Garimella KV, Maguire JR, Hartl C, Philippakis AA, del Angel G, Rivas MA, Hanna M, McKenna A, Fennell TJ, Kernytsky AM, Sivachenko AY, Cibulskis K, Gabriel SB, Altshuler D, Daly MJ. 2011. A framework for variation discovery and genotyping using next-generation DNA sequencing data. Nat Genet 43:491–8.

31. Van der Auwera GA, Carneiro MO, Hartl C, Poplin R, Del Angel G, Levy-Moonshine A, Jordan T, Shakir K, Roazen D, Thibault J, Banks E, Garimella KV, Altshuler D, Gabriel S, DePristo MA. 2013. From FastQ data to high confidence variant calls: the Genome Analysis Toolkit best practices pipeline. Curr Protoc Bioinformatics 43:11 10 1-11 10 33.

32. Cingolani P, Platts A, Wang le L, Coon M, Nguyen T, Wang L, Land SJ, Lu X, Ruden DM. 2012. A program for annotating and predicting the effects of single nucleotide polymorphisms, SnpEff: SNPs in the genome of *Drosophila melanogaste*r strain w1118; iso-2; iso-3. Fly (Austin) 6:80–92.

33. Cecil RE, Yoder-Himes DR. 2024. Examining the influence of environmental factors on *Acanthamoeba castellanii* and *Pseudomonas aeruginosa* in co-culture. PLoS One 19:e0305973.

34. Mojesky AA, Remold SK. 2020. Spatial structure maintains diversity of pyocin inhibition in household *Pseudomonas aeruginosa*. Proc Biol Sci 287:20201706.

35. Purdy-Gibson ME, France M, Hundley TC, Eid N, Remold SK. 2015. *Pseudomonas aeruginosa* in CF and non-CF homes is found predominantly in drains. J Cyst Fibros 14:341–6.

36. Remold SK, Brown CK, Farris JE, Hundley TC, Perpich JA, Purdy ME. 2011. Differential habitat use and niche partitioning by *Pseudomonas* species in human homes. Microb Ecol 62:505–17.

37. Hall CL, Lee VT. 2018. Cyclic-di-GMP regulation of virulence in bacterial pathogens. Wiley Interdiscip Rev RNA 9.

38. Coggan KA, Wolfgang MC. 2012. Global regulatory pathways and cross-talk control *Pseudomonas aeruginosa* environmental lifestyle and virulence phenotype. Curr Issues Mol Biol 14:47–70.

39. Hou L, Debru A, Chen Q, Bao Q, Li K. 2019. AmrZ Regulates Swarming Motility Through Cyclic di-GMP-Dependent Motility Inhibition and Controlling Pel Polysaccharide Production in *Pseudomonas aeruginosa* PA14. Front Microbiol 10:1847.

40. Romling U, Galperin MY, Gomelsky M. 2013. Cyclic di-GMP: the first 25 years of a universal bacterial second messenger. Microbiol Mol Biol Rev 77:1–52.

41. Ramsey DM, Wozniak DJ. 2005. Understanding the control of *Pseudomonas aeruginosa* alginate synthesis and the prospects for management of chronic infections in cystic fibrosis. Mol Microbiol 56:309–22.

42. Bernier SP, Ha DG, Khan W, Merritt JH, O’Toole GA. 2011. Modulation of *Pseudomonas aeruginosa* surface-associated group behaviors by individual amino acids through c-di-GMP signaling. Res Microbiol 162:680–8.

43. Palmer KL, Mashburn LM, Singh PK, Whiteley M. 2005. Cystic fibrosis sputum supports growth and cues key aspects of *Pseudomonas aeruginosa* physiology. J Bacteriol 187:5267–77.

44. Palmer KL, Aye LM, Whiteley M. 2007. Nutritional cues control *Pseudomonas aeruginosa* multicellular behavior in cystic fibrosis sputum. J Bacteriol 189:8079–87.

45. MJ A, LO M, I SC. 1991. Temperature profiles and alginate synthesis in mucoid and non mucoid variants of *Pseudomonas aeruginosa*. Letters in Applied Microbiology 12:244–248.

46. Westfall CS, Levin PA. 2017. Bacterial Cell Size: Multifactorial and Multifaceted. Annu Rev Microbiol 71:499–517.

47. Wang JD, Levin PA. 2009. Metabolism, cell growth and the bacterial cell cycle. Nat Rev Microbiol 7:822–7.

48. Price-Whelan A, Dietrich LE, Newman DK. 2007. Pyocyanin alters redox homeostasis and carbon flux through central metabolic pathways in *Pseudomonas aeruginosa* PA14. J Bacteriol 189:6372–81.

49. Eschbach M, Schreiber K, Trunk K, Buer J, Jahn D, Schobert M. 2004. Long-term anaerobic survival of the opportunistic pathogen *Pseudomonas aeruginosa* via pyruvate fermentation. J Bacteriol 186:4596–604.

50. Costa KC, Glasser NR, Conway SJ, Newman DK. 2017. Pyocyanin degradation by a tautomerizing demethylase inhibits *Pseudomonas aeruginosa* biofilms. Science 355:170–173.

51. Cox CD, Adams P. 1985. Siderophore activity of pyoverdin for *Pseudomonas aeruginosa*. Infect Immun 48:130–8.

52. Jeong GJ, Khan F, Khan S, Tabassum N, Mehta S, Kim YM. 2023. *Pseudomonas aeruginosa* virulence attenuation by inhibiting siderophore functions. Appl Microbiol Biotechnol 107:1019–1038.

53. Kuburich NA, Adhikari N, Hadwiger JA. 2016. *Acanthamoeba* and *Dictyostelium* Use Different Foraging Strategies. Protist 167:511–525.

54. Kumar A, Molli PR, Pakala SB, Bui Nguyen TM, Rayala SK, Kumar R. 2009. PAK thread from amoeba to mammals. J Cell Biochem 107:579–85.

55. Al-Quadan T, Price CT, Abu Kwaik Y. 2012. Exploitation of evolutionarily conserved amoeba and mammalian processes by *Legionella*. Trends Microbiol 20:299–306.

56. Siddiqui R, Khan NA. 2012. Biology and pathogenesis of Acanthamoeba. Parasit Vectors 5:6.

57. Guerin J, Bigot S, Schneider R, Buchanan SK, Jacob-Dubuisson F. 2017. Two-Partner Secretion: Combining Efficiency and Simplicity in the Secretion of Large Proteins for Bacteria-Host and Bacteria-Bacteria Interactions. Front Cell Infect Microbiol 7:148.

58. Kinoshita M, Namba K, Minamino T. 2021. A positive charge region of *Salmonella* FliI is required for ATPase formation and efficient flagellar protein export. Commun Biol 4:464.

59. Zhuang WY, Shapiro L. 1995. *Caulobacter* FliQ and FliR membrane proteins, required for flagellar biogenesis and cell division, belong to a family of virulence factor export proteins. J Bacteriol 177:343–56.

60. Qi X, Xu X, Li H, Pan Y, Katharine Kraco E, Zheng J, Lin M, Jiang X. 2022. fliA, flrB, and fliR regulate adhesion by controlling the expression of critical virulence genes in Vibrio harveyi. Gene 839:146726.

61. Flores-Bautista E, Cronick CL, Fersaca AR, Martinez-Nunez MA, Perez-Rueda E. 2018. Functional Prediction of Hypothetical Transcription Factors of Escherichia coli K-12 Based on Expression Data. Comput Struct Biotechnol J 16:157–166.

62. Lenski RE. 2017. Experimental evolution and the dynamics of adaptation and genome evolution in microbial populations. ISME J 11:2181–2194.

63. Ma L, Conover M, Lu H, Parsek MR, Bayles K, Wozniak DJ. 2009. Assembly and development of the Pseudomonas aeruginosa biofilm matrix. PLoS Pathog 5:e1000354.

64. Ryder C, Byrd M, Wozniak DJ. 2007. Role of polysaccharides in Pseudomonas aeruginosa biofilm development. Curr Opin Microbiol 10:644–8.

65. Toyofuku M, Roschitzki B, Riedel K, Eberl L. 2012. Identification of proteins associated with the Pseudomonas aeruginosa biofilm extracellular matrix. J Proteome Res 11:4906–15.

66. York A. 2018. Sharing the burden to build a matrix. Nat Rev Microbiol 16:454–455.

67. Leong W, Poh WH, Williams J, Lutz C, Hoque MM, Poh YH, Yee BYK, Chua C, Givskov M, Sanderson-Smith M, Rice SA, McDougald D. 2022. Adaptation to an Amoeba Host Leads to Pseudomonas aeruginosa Isolates with Attenuated Virulence. Appl Environ Microbiol 88:e0232221.

68. Davies EV, James CE, Brockhurst MA, Winstanley C. 2017. Evolutionary diversification of Pseudomonas aeruginosa in an artificial sputum model. BMC Microbiol 17:3.

69. Valentin JDP, Straub H, Pietsch F, Lemare M, Ahrens CH, Schreiber F, Webb JS, van der Mei HC, Ren Q. 2022. Role of the flagellar hook in the structural development and antibiotic tolerance of Pseudomonas aeruginosa biofilms. The ISME Journal 16:1176–1186.

70. Du B, Gu Y, Chen G, Wang G, Liu L. 2020. Flagellar motility mediates early-stage biofilm formation in oligotrophic aquatic environment. Ecotoxicology and Environmental Safety 194:110340.

71. Valentin JDP, Altenried S, Varadarajan AR, Ahrens CH, Schreiber F, Webb JS, van der Mei HC, Ren Q. 2023. Identification of Potential Antimicrobial Targets of Pseudomonas aeruginosa Biofilms through a Novel Screening Approach. Microbiol Spectr 11:e0309922.

72. Demirdjian S, Sanchez H, Hopkins D, Berwin B. 2019. Motility-Independent Formation of Antibiotic-Tolerant Pseudomonas aeruginosa Aggregates. Appl Environ Microbiol 85.

73. Klausen M, Heydorn A, Ragas P, Lambertsen L, Aaes-Jørgensen A, Molin S, Tolker-Nielsen T. 2003. Biofilm formation by Pseudomonas aeruginosa wild type, flagella and type IV pili mutants. Molecular Microbiology 48:1511–1524.

74. Nasher F, Wren BW. 2023. Flagellin O-linked glycans are required for the interactions between Campylobacter jejuni and Acanthamoebae castellanii. Microbiology (Reading) 169.

75. Preston TM, King CA. 1984. Binding Sites for Bacterial Flagella at the Surface of the Soil Amoeba Acanthamoeba. Microbiology 130:1449–1458.

76. Finkel SE, Kolter R. 1999. Evolution of microbial diversity during prolonged starvation. Proc Natl Acad Sci U S A 96:4023–7.

77. Zion S, Katz S, Hershberg R. 2024. Escherichia coli adaptation under prolonged resource exhaustion is characterized by extreme parallelism and frequent historical contingency. PLoS Genet 20:e1011333.

78. Behringer MG, Ho WC, Miller SF, Worthan SB, Cen Z, Stikeleather R, Lynch M. 2024. Trade-offs, trade-ups, and high mutational parallelism underlie microbial adaptation during extreme cycles of feast and famine. Curr Biol 34:1403–1413 e5.

