## Supplementary figures and images for "Long-term Culturing of *Pseudomonas aeruginosa* in Static, Minimal Nutrient Medium Results in Increased Pyocyanin Production, Reduced Biofilm Production, and Loss of Motility"

### Figure S1

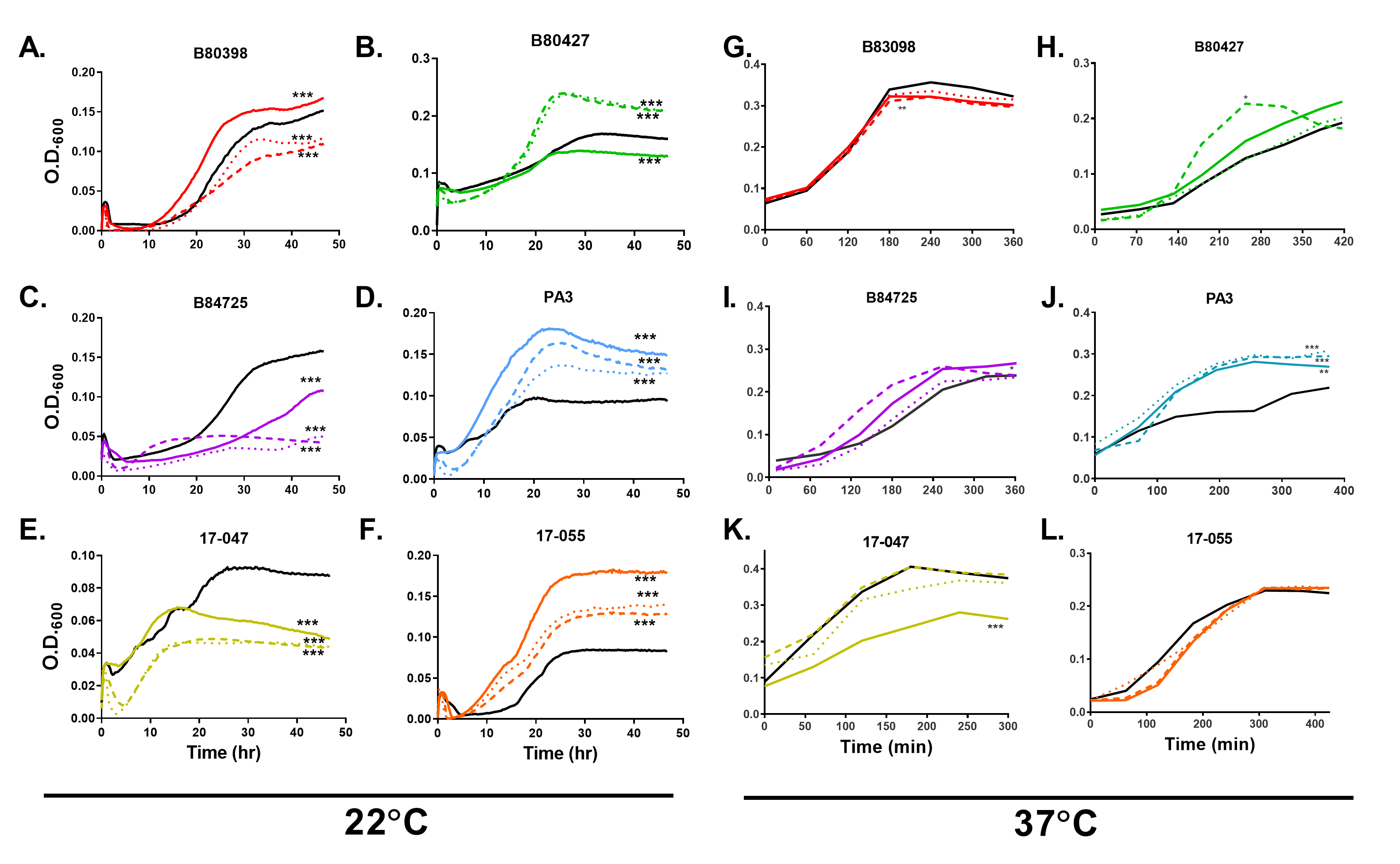

### Figure S2

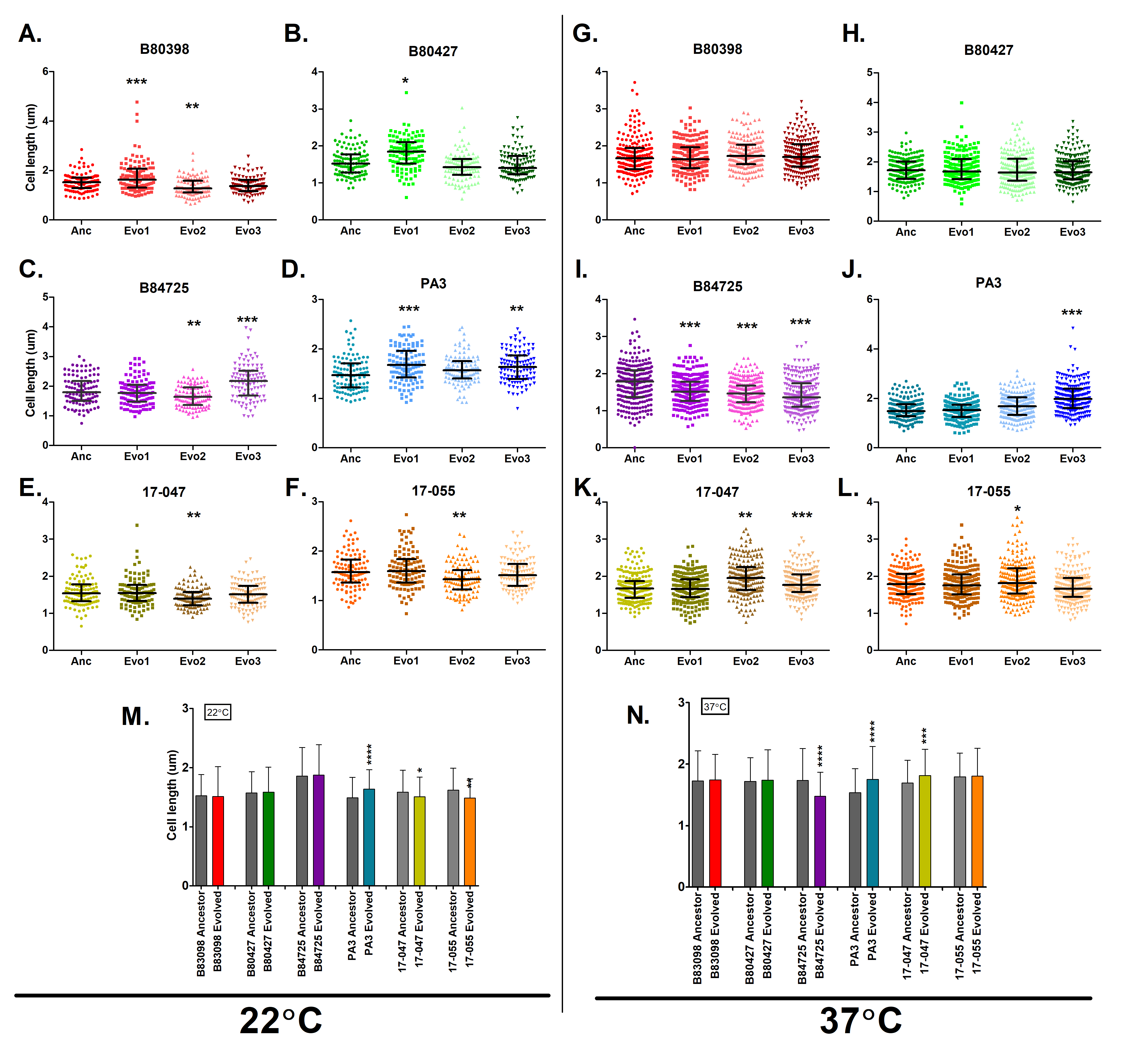

### Figure S3

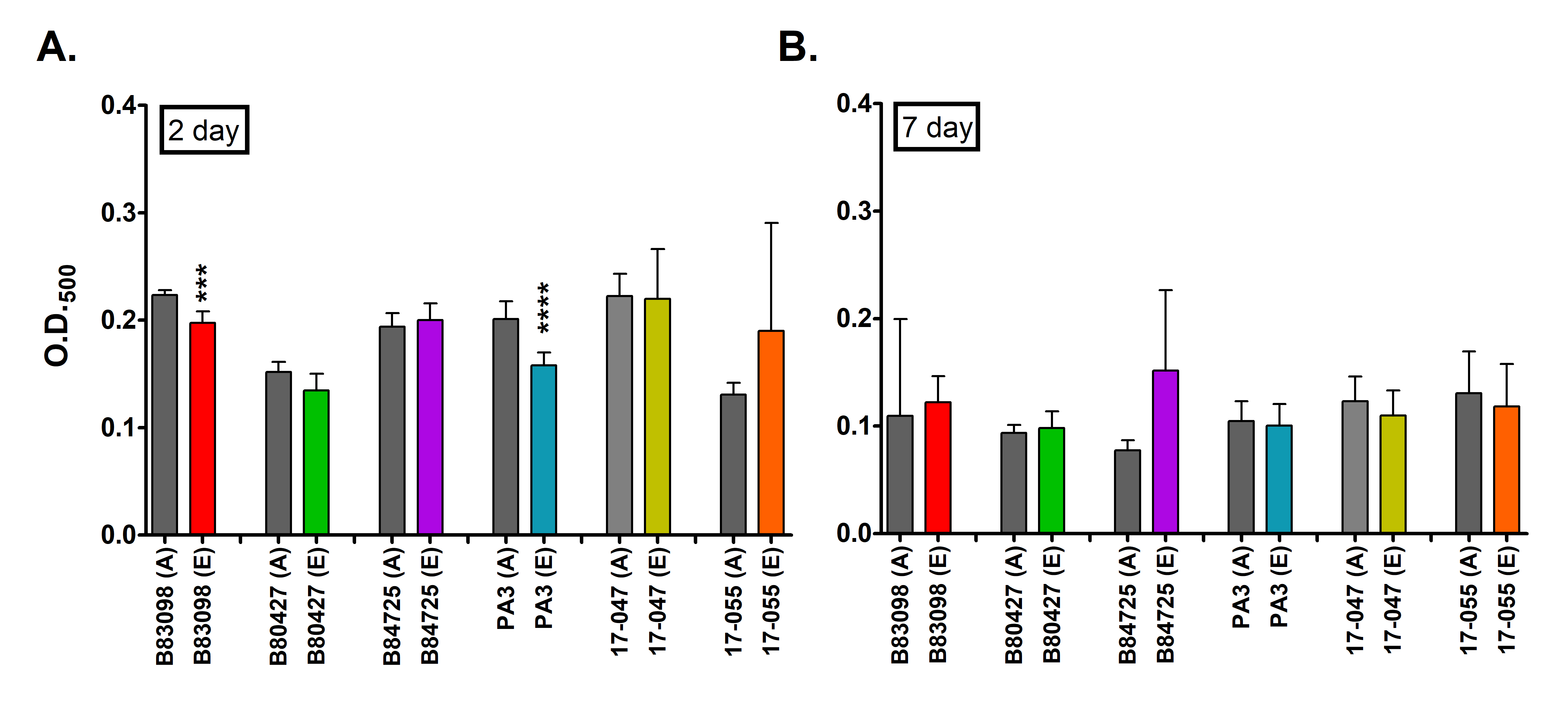

### Figure S4

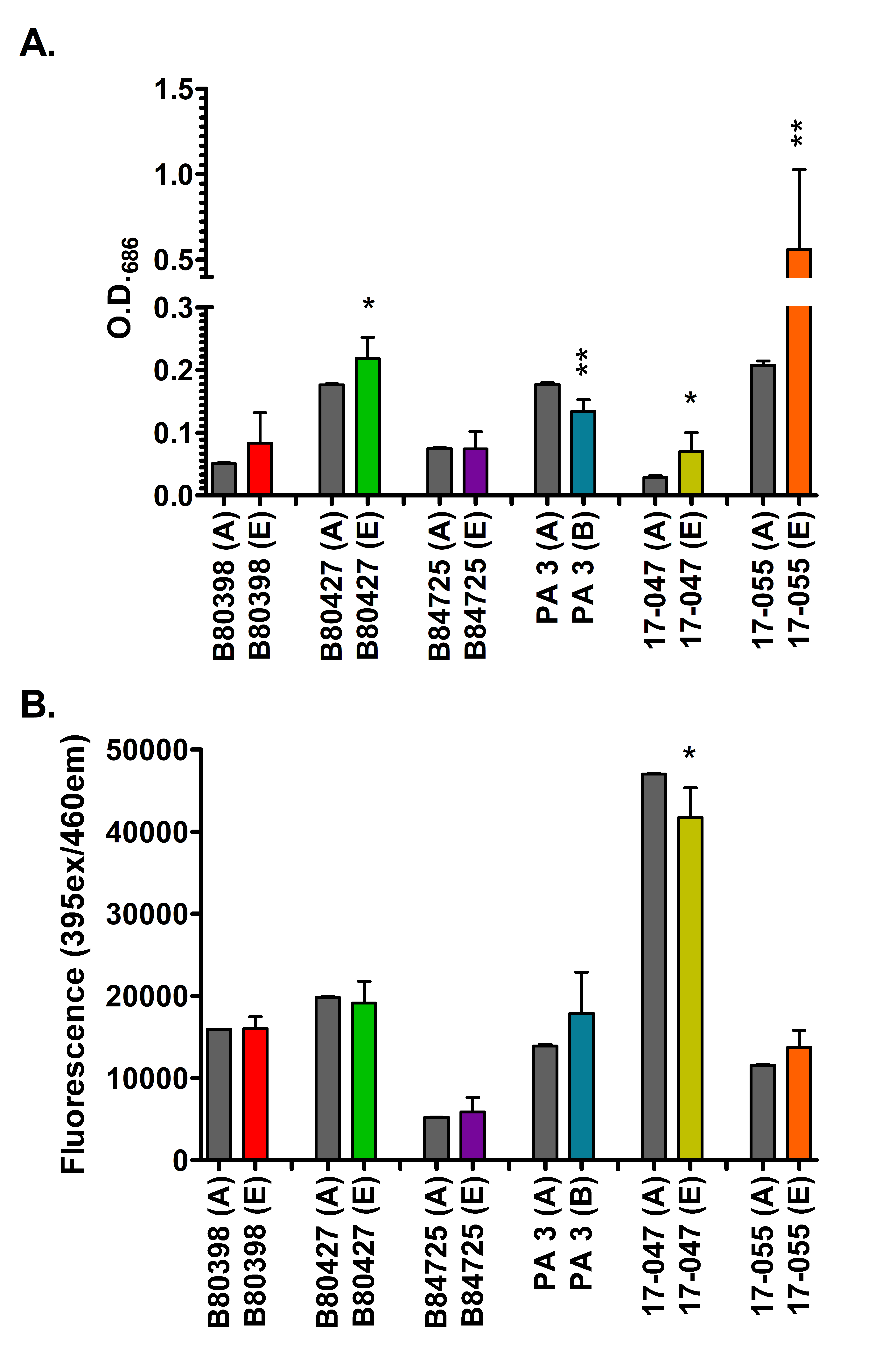

### Figure S5

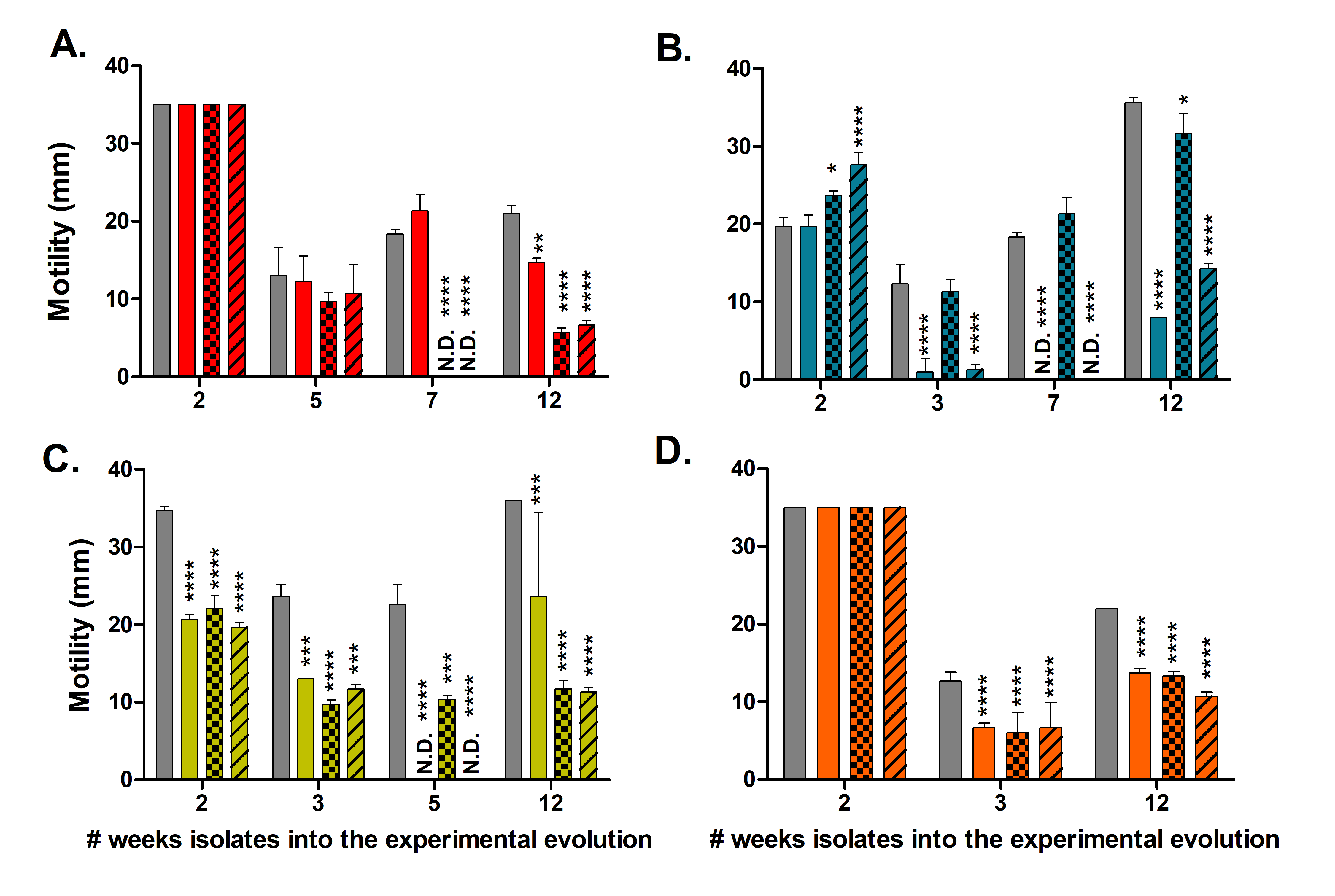

### Figure S6

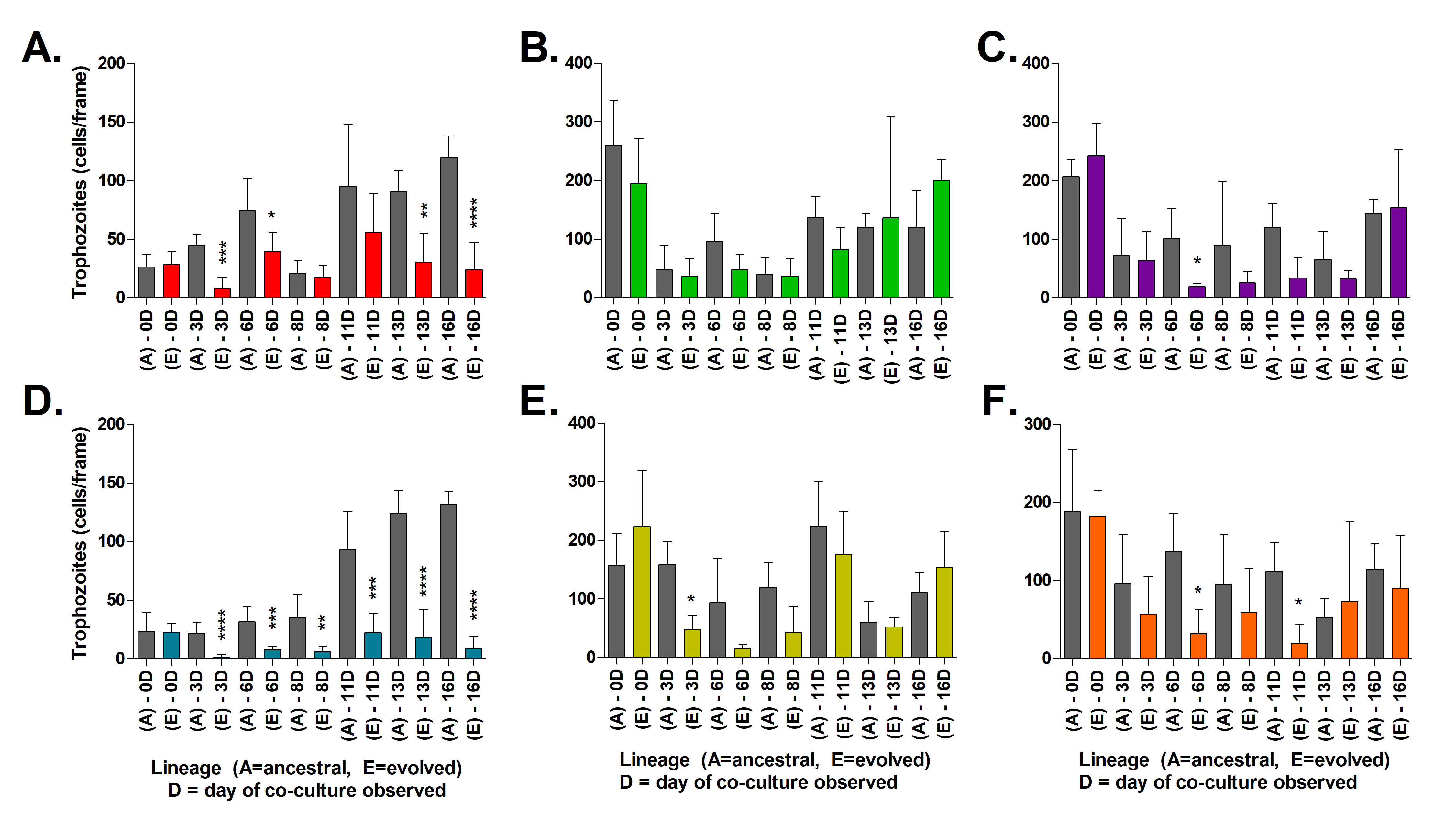
